# Synergistic effects of presynaptic and postsynaptic neurons give rise to three-stage short-term plasticity *in vivo*

**DOI:** 10.64898/2026.06.11.731703

**Authors:** Kai Lun Teh, Elena Dossi, Nathalie Rouach, Jérémie Sibille, Jens Kremkow

## Abstract

Short-term plasticity (STP) is the transient fluctuation of connection strength between two neurons depending on the recent history of neuronal activity. STP shapes neurotransmission over time and plays important roles in circuit computations. It is classically quantified *ex vivo* either at the synaptic level or at the level of spike transmission from the presynaptic neuron to the postsynaptic neuron. However, the exact relationship between the postsynaptic dendritic responses and spike transmission during STP still remains unclear *in vivo*. Here, we characterized together the STP of both postsynaptic dendritic responses, measured by the postsynaptic field potential (PFP), and spike transmission at the retinocollicular pathway of mice. We found mostly facilitating STP, where the second presynaptic spike occurring within 25 ms induces a larger PFP, and consequently a higher postsynaptic firing rate compared to responses from the first presynaptic spike. Both PFP and spike transmission exhibit short-term facilitation, but to a different degree, where the facilitation in the spike transmission is larger than the PFP. The PFP and spike transmission also exhibit facilitation of different decay time constants, indicating a nonlinear relationship between the two. Interestingly, a preceding postsynaptic spike can induce a similarly large but longer lasting facilitation on the spike transmission upon receiving a subsequent presynaptic input. However, STP of the PFP does not depend on the preceding postsynaptic spike, suggesting that this longer postsynaptic facilitation has a nonsynaptic origin. Overall, our results indicate that STP of the retinocollicular pathway exhibits three different stages: 1) a weak synaptic facilitation of postsynaptic dendritic responses, 2) a strong synaptic facilitation of spike transmission, and 3) a longer lasting nonsynaptic facilitation of spike transmission. Using a computational model, we show that the second STP stage is a direct inheritance from the first STP stage, whereas two opposing nonsynaptic mechanisms with different time constants are needed for the emergence of the third STP stage. These findings provide direct evidence that synaptic and nonsynaptic STPs coexist *in vivo*, paving the way for large-scale measurement of these STPs and offering a means to monitor the transmission of information in neural circuits of behaving animals.

## INTRODUCTION

Short-term plasticity (STP) plays a critical role in circuit dynamics and information transfer (Abbott and Regehr, 2004). It is the transient change of information transmission capacity at either the synaptic or neuronal level, and can be broadly classified as synaptic and nonsynaptic forms (Kern and Chao, 2023; Sánchez-Aguilera et al., 2014). Synapses act as gatekeepers for the transmission of information from one neuron to another, where they undergo history-dependent changes. This alters their transmission capacity (Abbott and Regehr, 2004; Fortune and Rose, 2001; Rotman et al., 2011), and thereby dynamically shapes information flow within neural circuits. These transient modifications form the basis of synaptic STP, which has been extensively studied *ex vivo* over the past few decades. Synaptic STP can be viewed as a temporary cache of network state, allowing neural circuits to rapidly access, retrieve, and compute relevant information (Del Gaudio et al., 2025; Kozachkov et al., 2022; Mongillo et al., 2008; Seeholzer et al., 2019; Taher et al., 2020). However, how synaptic STP regulates the postsynaptic spiking *in vivo* remains unclear.

Besides synapses, nonsynaptic components are also indispensable in the dynamical transmission of information (Ghanbari et al., 2020; Kern and Chao, 2023). Nonsynaptic STP primarily regulates intrinsic neuronal excitability, thereby modulating the input-output function of a neuron over short timescales (Egorov et al., 2002; Mahon et al., 2003; Sánchez-Aguilera et al., 2014), which has been implicated in diverse brain functions (Deng et al., 2024; Fujisawa et al., 2008; Kern and Chao, 2023; Motanis et al., 2018; Pellizzari et al., 2023; Zhang and Sillar, 2012). Yet the nonsynaptic STP *in vivo* is poorly understood and is challenging to infer solely from extracellular spikes (Ghanbari et al., 2020). In short, synaptic STP regulates the synaptic input strength from the presynaptic neuron to the postsynaptic neuron, whereas nonsynaptic STP governs the excitability of a neuron. Together, they orchestrate the dynamics of neural information transmission, forming the foundation of information processing in the brain (Kern and Chao, 2023). Although both synaptic and nonsynaptic STPs have been extensively investigated *in vitro* and *ex vivo* (Brody and Yue, 2000; Deng et al., 2024; Egorov et al., 2002; Martinetti et al., 2022; Sánchez-Aguilera et al., 2014; Xu and Wu, 2005), largely in isolation, their forms and interactions *in vivo* remain unclear, mainly due to technical constraints.

*In vivo* studies using intracellular recordings measured synaptic STP at thalamocortical synapses (Boudreau and Ferster, 2005; Chung et al., 2002; Jia et al., 2006). Other *in vivo* studies using extracellular recordings with probes inserted across cortical layers estimated the postsynaptic current sink using current source density, and thus indirectly measured synaptic STP (Bereshpolova et al., 2019; Swadlow et al., 2002). However, these studies mainly focused on the synaptic STP, leaving its relationships with postsynaptic spiking and nonsynaptic STP unexplored. Besides, these approaches often require topographic alignment between two different regions in order to capture connected neuron pairs, which is technically challenging and low-yield. Extracellular recordings with high-density electrodes have the advantage of measuring the neuronal spiking activities in large scale, and studies have shown that the synaptic properties and STP can be estimated by using the cross-correlation of the spike trains between the presynaptic and postsynaptic neurons (English et al., 2017; Fujisawa et al., 2008; Ghanbari et al., 2017). Nonetheless, these methods do not provide direct access to the underlying synaptic responses, leaving ambiguities between synaptic and nonsynaptic components, and thus isolating synaptic and nonsynaptic STPs remains challenging with these techniques. Therefore, how synaptic and nonsynaptic STPs coordinate *in vivo* is still unclear. More precisely, how the STP of synaptic inputs translates into the STP of somatic spikes in the postsynaptic neuron and to what extent the postsynaptic spiking is independent of the synaptic components are still perplexing. In other words, what are the relationships between the synaptic and nonsynaptic STPs *in vivo* and how do they coordinate in the brain? Can synaptic and nonsynaptic STPs be disentangled using extracellular recordings?

Modern electrophysiological techniques enable simultaneous measurement of synaptic and nonsynaptic components (Sibille et al., 2022a, 2024), providing a holistic view of local circuit dynamics. With these, it is possible to measure both postsynaptic dendritic responses and subsequent somatic spikes extracellularly in a living brain, offering a way to explore the relationships between the synaptic responses and the evoked spikes, as well as their corresponding STP. Here, we demonstrate the feasibility in determining the relationships between the synaptic and nonsynaptic STPs *in vivo* by simultaneously measuring the postsynaptic responses and both presynaptic and postsynaptic spiking activity (Figs. 1A-B). We found that the STP *in vivo* exhibits at least three different stages, where the synaptic responses and neuronal spiking can be coupled or decoupled, giving rise to diverse STPs with different timescales and decay time constants. Using Tsodyks-Markram synapses and a leaky integrate-and-fire (LIF) neuron model, we propose a mechanism that can account for the three observed STP stages together. These findings reinforce the classical view that STP occurs in multiple stages, while also extending this view by demonstrating that these STP phases synergistically interact *in vivo* as they can be directly detected using existing recording techniques. This study lays the groundwork for large-scale recording and monitoring of both synaptic and nonsynaptic STPs, taking a step toward a better understanding of circuit dynamics over different, and potentially longer, timescales in behaving animals (see Siegle and Steinmetz, 2026).

**Figure 1:**
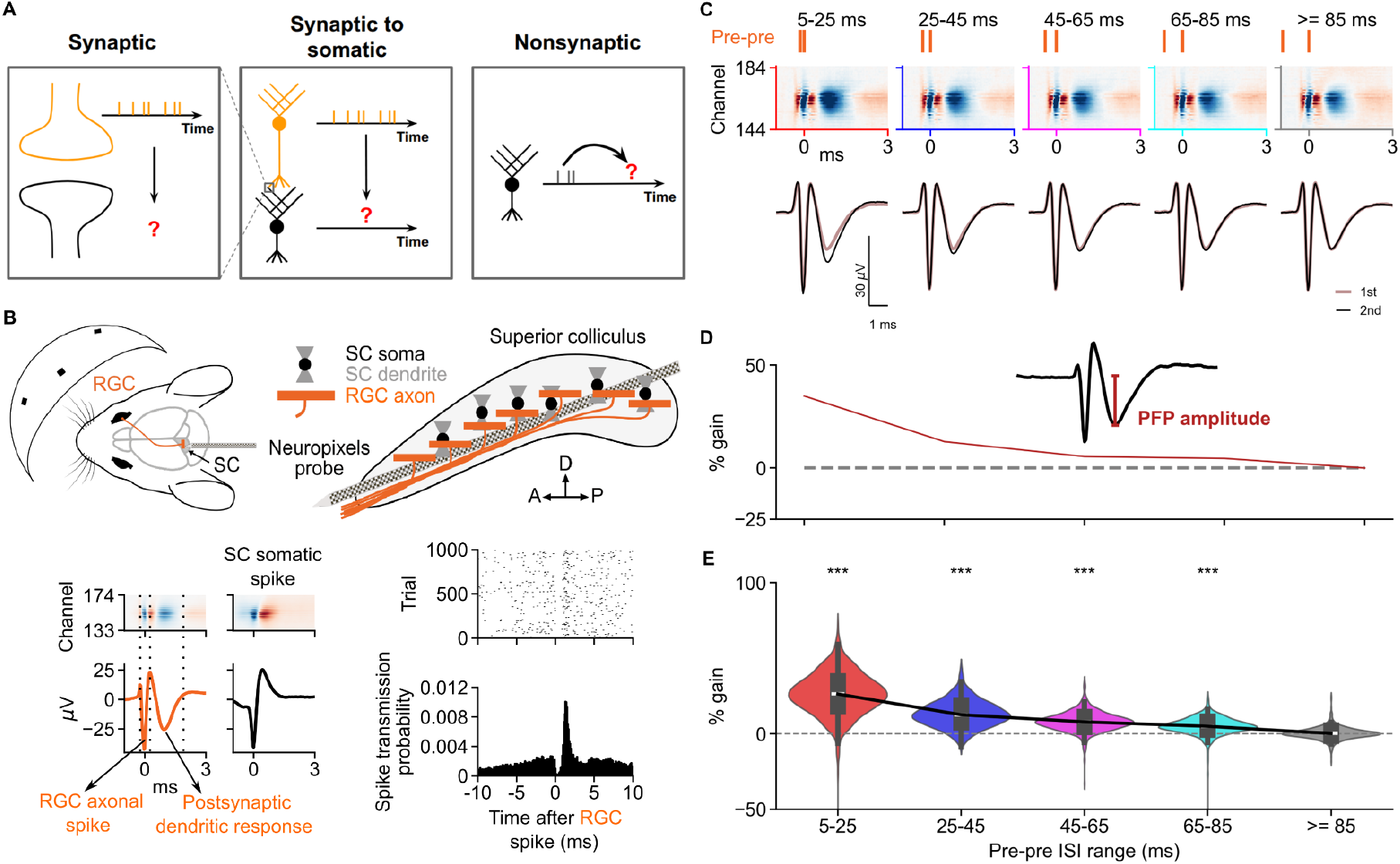
The PFP in SC reliably facilitates at short ISIs. (A) Three different synaptic and nonsynaptic STP stages can be measured extracellularly *in vivo* using existing electrophysiological techniques. (B) Top: scheme showing the experimental setup (adapted from Sibille et al., 2022a). Visual stimuli were presented to a mouse (left) with a Neuropixels probe tangentially inserted in the superficial layers of the contralateral SC (right). Bottom left: multichannel waveform and the corresponding one-dimensional waveform for a connected pair of an RGC (left) and an SC neuron (right). Bottom right: example raster trials of the postsynaptic SC neuron aligned to the spike times of the presynaptic RGC (top) and the corresponding cross-correlogram (bottom). (C) Top: scheme showing different pre-pre ISI ranges. Middle: multichannel waveforms of an RGC averaged over the second spikes of different pre-pre ISIs. Bottom: one-dimensional waveforms obtained by averaging across 11 channels (±5 channels from the best channel defined by Kilosort) of the multichannel waveforms. (D) The corresponding percentage gain in PFP amplitude of the second relative to the first (baseline) RGC spikes for the waveforms in C, where the PFP induced by the second RGC spike has a larger amplitude compared to the PFP induced by the first RGC spike for shorter ISIs. Inset indicates a typical recorded axonal waveform that has characteristic triphasic components (double negative peaks) with PFP amplitude labeled. (E) The distributions of percentage gain in PFP amplitude for different ISI ranges, where the ISI range of 5-25 ms has the largest gain. The black line indicates the median, the box inside the violin plot represents the Q1 to Q3 of the percentage amplitude gains (Q1: 17.9%, 7.3%, 4.3%, 2.55%, -1.62%; median: 26.26%, 12.49%, 7.79%, 4.86%, 0.12%; Q3: 34.77%, 18.64%, 11.23%, 8.18%, 2.13%; *p* = 9.63 x 10^-79^, 3.34 x 10^-76^, 5.9 x 10^-68^, 4.32 x 10^-56^, 0.333 for ISI ranges of 5-25, 25-45, 45-65, 65-85, and ≥85 ms, two-sided Wilcoxon signed-rank test for first PFP vs. second PFP, *n* = 498 RGCs, *n* = 27 experiments, *n* = 24 mice).

## RESULTS

### Characterizing the synaptic STP *in vivo*

We first examined the effects of paired presynaptic spikes on the postsynaptic responses, as classically studied *ex vivo* (Cho et al., 2011; Creager et al., 1980; Dobrunz et al., 1997; Yang and Xu-Friedman, 2008). In our previous study, we demonstrated the feasibility of simultaneously recording the postsynaptic signals of the retinocollicular synapses and the postsynaptic neuronal spikes in the superior colliculus (SC) (Sibille et al., 2022a). We now employed this approach to characterize the STP in the retinocollicular pathway. In these experiments, a Neuropixels probe was tangentially inserted into the visual layers of SC to record the extracellular activities while visual stimuli were presented to the animals to induce visual responses (Fig. 1B). The spikes of retinal ganglion cell (RGC) axons and SC neurons were distinguished based on their distinct waveforms (Fig. 1B, bottom left; Gehr et al., 2023; Sibille et al., 2022a). With this approach, we measured the axonal spikes of single RGCs and their corresponding postsynaptic responses, termed postsynaptic field potential (PFP), evoked in the SC by the RGC spike (Fig. 1C-D). More specifically, the PFP is an extracellular signal corresponding to the net excitation elicited by a single RGC axon onto postsynaptic dendritic sites of multiple SC neurons (Sauvé et al., 1995; Shein-Idelson et al., 2017; Sibille et al., 2022a). It can be thought of as the cumulative synchronous extracellular signal from all intracellular excitatory postsynaptic potentials (EPSP) induced by a single presynaptic RGC input spike.

To characterize the synaptic transmission properties of the retinocollicular pathway, we analyzed the percentage amplitude change of the average PFP induced by the second RGC spikes compared to the first RGC spikes within five ranges of presynaptic-presynaptic (pre-pre) interspike interval (ISI), namely, 5-25, 25-45, 45-65, 65-85, and ≥85 ms. We found that the shorter the pre-pre ISI, the larger the PFP amplitude evoked by the second RGC spike compared to the PFP amplitude evoked by the first RGC spike (Figs. 1C-E). However, a previous *ex vivo* study on the superficial SC of hamster SC slices observed short-term depression instead of facilitation by using the paired-pulse protocol (Balmer and Pallas, 2015). To strengthen our *in vivo* observations, we performed experiments in acute mouse SC slices (Figs. S1A-B), and observed similar facilitating effects in the paired-pulse protocol (Fig. S1C) as well as repetitive stimulations (Fig. S1D). In our *in vivo* observations, the peak-to-peak axonal action potential (AP) amplitude remained largely unchanged between the first and second RGC spike (Fig. S2A), thereby serving as a control.

To make sure that there are no confounding effects on the PFP signals by the postsynaptic SC neuronal spikes (Sauvé et al., 1995), we compared different variables of the average RGC waveforms with and without tailgating postsynaptic SC spikes (Fig. S3). A new average RGC waveform was estimated without SC spikes, computed by removing the RGC spikes that have any detectable SC spikes occurring within the lag window [-3, 6] ms before averaging. No obvious difference in the PFP facilitation was found between the RGC waveforms with and without tailgating SC spikes for all variables quantified (Fig. S3). To further make sure that the postsynaptic SC spiking signals do not contribute to the PFP amplitude, we applied 2.5 mM muscimol, a potent GABA_A_ agonist, in a subset of experiments to inhibit the SC neuronal spiking. The visual stimuli were presented 20 minutes after the muscimol application. Comparison between the control condition and muscimol application in SC showed no obvious difference in the PFP facilitation for all waveform variables quantified (Fig. S4). Together, the results of these two controls further support the observation that PFP amplitude facilitates at shorter ISIs (Fig. 1E).

Since RGCs make strong and specific functional connections onto SC neurons (Sibille et al., 2022a) of both excitatory and inhibitory types (Gehr et al., 2023), the RGC axonal boutons might be overactivated and thus giving a small PFP facilitation (Fig. 1E) due to presynaptic vesicle depletion, where further RGC spikes might lead to depression instead. To determine whether the RGC axonal boutons are overactivated, we analyzed the PFP amplitudes induced by multiple consecutive RGC spikes with regular ISI ranges of 5-25 ms (Fig. S2B, top). We found that the PFP amplitude keeps increasing for at least three (Fig. S2B, middle) and four consecutive spikes (Fig. S2B, right) with similar ISIs in comparison to the PFP induced by the first RGC spike, suggesting that the observed facilitation can continue over multiple consecutive spikes. Again, similar observations were found in the SC slice recordings (Fig. S1D), indicating that the synaptic transmission facilitation at the retinocollicular synapses is not saturated at least for four consecutive spikes, in both *in vivo* and *ex vivo* settings. In short, the larger PFP amplitude at shorter ISIs, together with the increasing PFP amplitude in several consecutive spikes, indicate that the retinocollicular synapses mainly undergo a small, but reliable, short-term facilitation.

### Spike transmission probability *in vivo*

We have shown that short pre-pre ISIs can induce facilitation at the PFP level, but does this PFP facilitation translate into a higher spiking output of the postsynaptic SC neurons? First, the potential connection between an RGC-SC neuron pair (Fig. 2A) was determined by quantifying the average spike transmission probability from the spike train cross-correlogram (CCG; Fig. 2B; English et al., 2017; Sibille et al., 2022a; Stark and Abeles, 2009). More specifically, the average spike transmission probability was computed by subtracting the baseline (Fig. 2B, light green line) from the CCG (baseline corrected CCG) and summing the positive peak area between the lags of 0.8 ms and 2.8 ms. To investigate the effects of the pre-pre ISI on the spike transmission probability, we split the RGC spike pairs into five pre-pre ISI ranges as before, and quantified the spike transmission probability resulting from the pre-pre ISI grouped CCGs of connected RGC-SC neuron pairs (Fig. 2C, D; see Methods) separately for the first and second RGC spikes. The gain in the spike transmission probability was computed by subtracting the spike transmission probability of the first RGC spikes (Fig. 2D, top) from the second RGC spikes (Fig. 2D, bottom) for each pre-pre ISI range. This gain was then normalized to the average pre-pre ISI spike transmission probability to obtain the percentage gain in the spike transmission probability. The spike transmission probability of the average pre-pre ISI, instead of the first RGC spike, was used because some RGC-SC neuron pairs have little to no spike transmission for the first RGC spikes, which will result in inaccurate estimation of gain. Despite using the average pre-pre ISI spike transmission probability as the baseline for normalization underestimates the spike transmission gain in comparison to using the first RGC spike, it gives more consistent gain estimation.

**Figure 2:**
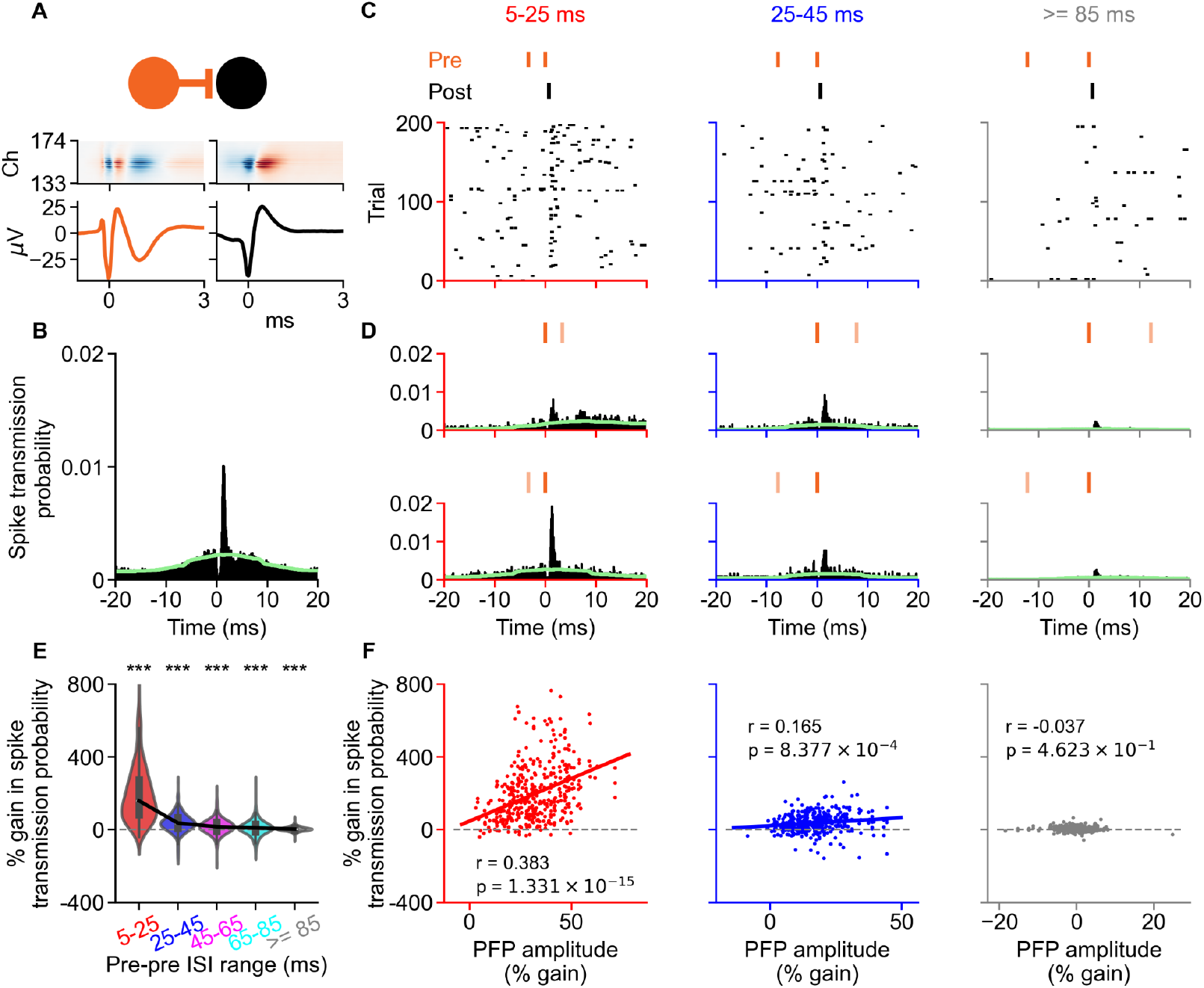
The PFP facilitation nonlinearly enhances the gain in spike transmission probability. (A) Scheme showing the presynaptic RGC (orange) and its postsynaptic SC partner (black) with their corresponding one-dimensional and two-dimensional waveforms. (B) CCG showing the spike transmission probability of a connected pair averaged over all presynaptic RGC spikes. The light green line indicates the CCG baseline used for computing the baseline corrected CCG. (C) Raster plots showing the postsynaptic SC activities upon receiving the second presynaptic input with ISIs of 5-25, 25-45, and ≥85 ms. (D) CCG showing the spike transmission probability of a connected pair averaged over the first (top) and second (bottom) RGC spikes of corresponding ISI ranges in C. (E) The shortest ISI range (5-25 ms) has the largest percentage gain in spike transmission probability, which decreases almost back to baseline at 45-65 ms. The black line indicates the median, the box inside the violin plot represents the Q1 to Q3 of the data (Q1: 80.62%, 7.95%, -7.44%, -12.28%, -3.35%; median: 158.49%, 35.61%, 15.79%, 8.79%, 3.26%; Q3: 274.67%, 65.53%, 39.21%, 30.39%, 9.89%; *p* = 3.38 x 10^-67^, 6.36 x 10^-40^, 4.69 x 10^-19^, 5.34 x 10^-8^, 1.44 x 10^-8^ for ISI ranges of 5-25, 25-45, 45-65, 65-85, and ≥85 ms, two-sided Wilcoxon signed-rank test for first spike vs. second spike, *n* = 406 pairs, *n* = 224 RGCs, *n* = 21 experiments, *n* = 20 mice). (F) The percentage gain in the spike transmission probability is positively correlated to the facilitation of the PFP amplitude. The 5-25 ms ISI range has the highest correlation (left) compared to other ISI ranges (middle and right).

Similar to the PFP amplitude, the percentage gain in the spike transmission is higher at shorter pre-pre ISIs (Fig. 2E). The percentage gain in the spike transmission probability has a strong positive correlation to the facilitation of the PFP amplitude at 5-25 ms ISI range (Fig. 2F, left; *r* = 0.383, *p* = 1.33 x 10^-15^, *n* = 406 pairs), indicating that the facilitation of PFP amplitude can enhance the spike transmission probability for shorter ISIs. Similarly, the second largest ISI range of 25-45 ms has also a small, but significant positive correlation between the percentage gain in the spike transmission probability and the STP of the PFP amplitude (Fig. 2F, middle; *r* = 0.165, *p* = 8.38 x 10^-4^, *n* = 406 pairs). This suggests a presynaptic origin for the facilitation of postsynaptic spiking during the pre-pre ISI condition.

The median percentage gain in PFP amplitude of the second RGC spike is around 26% for the 5-25 ms, which is a relatively small increase when compared to the median percentage gain of ∼158% in spike transmission probability. When fitted to an exponential decay function of the form 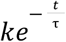, where *k* is the initial value of the exponential decay function at *t* = 0 ms and τ is the decay time constant, the pre-pre facilitation of PFP amplitudes has longer decay time constants and smaller initial values (Fig. S5A-C) in comparison to the pre-pre facilitation of spike transmissions (Fig. S5D-F). The median of spike transmission facilitation decays with a time constant of 14.49 ms (Fig. S5F), which is shorter than the decay time constant of the median PFP facilitation (31.78 ms; Fig. S5C). Together with the decreasing slopes of the fitted relationships across the pre-pre ISI ranges (4.705 vs. 0.877 for ISI ranges of 5-25 vs. 25-45 ms; Fig. 2F), these observations indicate a nonlinear relationship between the PFP facilitation and the spike transmission facilitation over the pre-pre ISI ranges. To sum up, spike transmission in the pre-pre ISI condition is facilitating and is correlated to the PFP facilitation, although they have different decay time constants and the large gain in spike transmission might not be entirely attributed to the small PFP facilitation.

### Nonsynaptic short-term facilitation *in vivo*

Next, we examined how the preceding postsynaptic spike timing influences the STP upon the arrival of a presynaptic spike (English et al., 2017). This allowed us to determine whether a previous activation of a postsynaptic SC neuron could have an effect on its responses to the next presynaptic input. To answer this question, we investigated the effects of the interval from the last postsynaptic spike to the next presynaptic spike, termed postsynaptic-presynaptic (post-pre) ISI, on the PFP and spike transmission of each connected pair. To ensure that there is no potential effect from the pre-pre ISI condition, we applied an exclusion criterion that a post-pre ISI condition is valid only if the presynaptic spike has a dead time of at least 85 ms. Similar to the pre-pre ISIs, the post-pre ISIs were categorized into the five same ISI ranges.

In the post-pre ISI condition, there is a gap roughly equal to the smallest post-pre ISI of each group, which can be seen from the raster plots (Fig. 3A) and the corresponding CCGs (Fig. 3B). This ensures that for each ISI range, the connected postsynaptic neuron is silent for at least the smallest post-pre ISI. Our results show that the shorter the post-pre ISI, the higher the spike transmission probability upon the next presynaptic input (Fig. 3C). Moreover, the spike transmission facilitation of the post-pre ISI (Fig. 3C) lasts longer than the pre-pre ISI (Fig. 2E), where it is more steady and does not exhibit a simple exponential decay trend.

**Figure 3:**
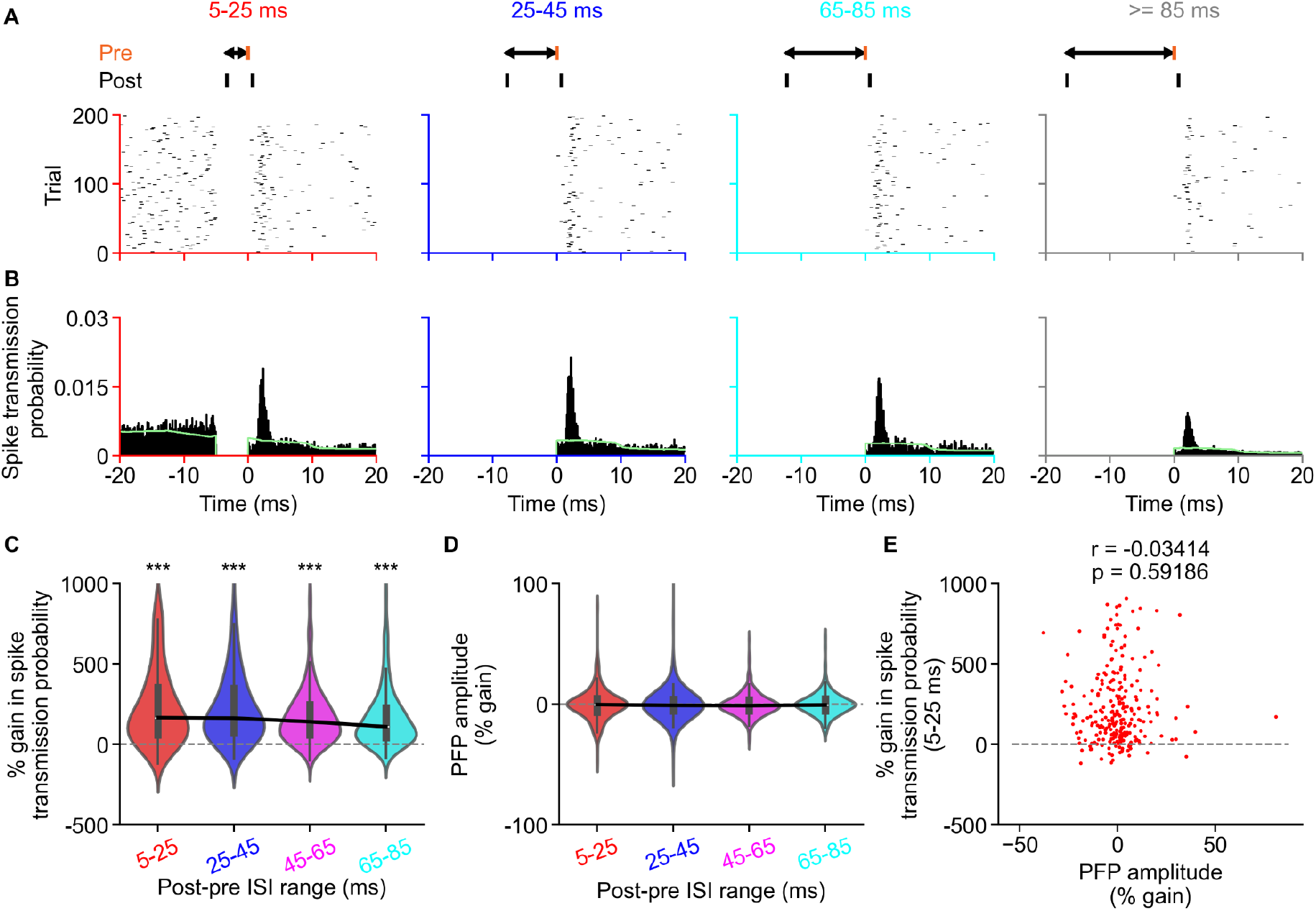
Postsynaptic spiking induces facilitation of spike transmission independent of PFP facilitation. (A) Raster plots showing the postsynaptic activities upon receiving the presynaptic inputs of different post-pre ISIs. (B) CCGs showing the spike transmission probability of a connected pair computed with corresponding post-pre ISIs in A. The light green line indicates the CCG baseline. (C) The percentage gain in spike transmission probability normalized to the average post-pre ISI spike transmission probability against different post-pre ISI ranges. The black line indicates the median, the box inside the violin plot represents the Q1 to Q3 of the data (Q1: 57.46%, 72.38%, 57.63%, 42.23%; median: 166.11%, 163.3%, 140.82%, 109.71%; Q3: 352.35%, 347.1%, 247.26%, 226.88%; *p* = 4.61 x 10^-37^, 3.72 x 10^-40^, 2.56 x 10^-40^, 9.48 x 10^-40^ for ISI ranges of 5-25, 25-45, 45-65, and 65-85 ms, two-sided Wilcoxon signed-rank test for test ISI range vs. baseline ISI range of ≥85 ms, *n* = 249 pairs, *n* = 168 RGCs, *n* = 19 experiments, *n* = 18 mice). (D) No facilitation in the PFP for the post-pre ISIs. The black line indicates the median, the box inside the violin plot represents the Q1 to Q3 of the data (Q1: -6.74%, -5.53%, -5.4%, -5.6%; median: -0.36%, -1.05%, -1.29%, -0.65%; Q3: 4.53%, 4.08%, 3.52%, 4.41%; *p* = 0.411, 0.239, 0.056, 0.441 for ISI ranges of 5-25, 25-45, 45-65, and 65-85 ms, two-sided Wilcoxon signed-rank test, *n* = 249 pairs, *n* = 168 RGCs, *n* = 19 experiments, *n* = 18 mice). (E) There is no correlation between the percentage gain in PFP amplitude and the percentage gain in spike transmission probability for the post-pre ISI (5-25 ms) condition.

To our surprise, there was no facilitation observed in the PFP amplitude computed with the post-pre ISI condition (Fig. 3D), suggesting that the presynaptic input is likely not responsible for the increase in spike transmission under this condition. There is indeed no apparent correlation between the percentage gain in the post-pre ISI spike transmission probability and the percentage facilitation of the PFP amplitude at 5-25 ms (Fig. 3E; *r* = -0.034, *p* = 0.592, *n* = 249 pairs). However, there are strong negative correlations between the percentage gain in the post-pre ISI spike transmission probability and the average post-pre ISI spike transmission probability for all ISI ranges (Fig. S6). This suggests that the RGC-SC neuron pairs that have a strong connection are less likely to facilitate from the preceding postsynaptic spiking. In short, SC neurons are more excitable shortly after their previous spike (Fig. 3C), even though there is no apparent increase in the postsynaptic responses (Fig. 3D), and this type of facilitation is more prominent in weaker connections (Fig. S6).

### Short-term facilitation is largely independent of converging input rates

The pre-pre facilitations of the PFP and spike transmission are not limited to the paired spikes stemming from the same presynaptic RGC. The paired spikes from different convergent RGCs (Fig. S7A), where the number of convergent RGCs to a postsynaptic SC neuron follows a long-tailed distribution (Fig. S7B), also exhibit small but significant facilitation for both PFP and spike transmission, with the PFP facilitation (Fig. S7C-G) being less prominent than the spike transmission facilitation (Fig. S7H-L). Further categorization of the convergent RGCs into weak (average spike transmission probability < 0.02) and strong (average spike transmission probability ≥ 0.02) connections reveal that the spike transmission of a convergent pair facilitates more when the second presynaptic spike originates from an RGC with a strong connection to the postsynaptic SC neuron, regardless of the connection strength of the first RGC (Figs. S7I-L). In other words, if the second RGC has sufficiently high efficacy, then a prior spike from another connected RGC, regardless of its strength, will always lead to a facilitation of the spike transmission in the second RGC.

Given these small convergent facilitations, a potential confounding factor to the apparent pre-pre facilitation from the same presynaptic RGC is the co-firing of the convergent presynaptic RGCs. To further examine how the convergent input rate affects the STP, we categorized the valid spike trials in the pre-pre and post-pre conditions of the same ISI groups into five different input-rate groups, namely 0-10, 10-20, 20-30, 30-40, and ≥ 40 Hz. These converging input rates were defined by the number of spikes of other presynaptic RGCs within a 100 ms window prior to the current valid RGC spike of the pre-pre (Fig. S8A) and post-pre (Fig. S8B) ISI conditions.

The raw spike transmission gain at 5-25 ms pre-pre ISI group is always higher than the spike transmission gain in other ISI groups, where the spike transmission gain is almost back to 0 at 25-45 ms group for all input-rate groups (Fig. S8C). This indicates that the input rate does not modify the general trend of the pre-pre ISI facilitation. That is, within a rate group, the shorter the ISI, the higher the spike transmission gain. However, the higher input rates seem to slightly enhance the raw spike transmission gain of the 5-25 ms pre-pre ISI group (Fig. S8D), although the effect size is small (Cohen’s d = 0.381). On the other hand, the raw spike transmission gain of the post-pre ISI condition appears to be fairly independent of the input rates (Fig. S8E-F). The distributions of the spike transmission gains for 5-25 ms ISI groups are largely overlapped across different input-rate groups for both pre-pre (Fig. S8G) and post-pre (Fig. S8H) ISI conditions.

In short, higher input rates slightly enhance the raw spike transmission gain for pre-pre but not post-pre ISI conditions, and have little to no effect on the facilitating trend across ISI groups in both conditions. This implies that the pre-pre and post-pre ISI facilitations are likely not influenced by other presynaptic inputs to the same postsynaptic SC neurons.

### Modelling the three stages of short-term plasticity *in vivo*

We have shown that the retinocollicular pathway exhibits short-term facilitation differentially at the postsynaptic response level and at the spiking level depending either on the presynaptic or the postsynaptic spiking history. We classified these facilitations into three different stages (Fig. 4A): 1) the presynaptic-induced postsynaptic response, 2) the presynaptic-induced postsynaptic spiking, and 3) the postsynaptic-dependent postsynaptic spiking. To determine the potential mechanisms underlying the observed STPs, we used the leaky integrate-and-fire (LIF) neuron model with two optional additional mechanisms, spike adaptation current (Brette and Gerstner, 2005) and dynamic threshold (Horn and Opher, 2000), to reproduce our experimental observations. The full model consists of five Poisson presynaptic neurons with Tsodyks-Markram synapses (Mongillo et al., 2008; Tsodyks and Markram, 1997) and one adaptive LIF postsynaptic neuron (Brette and Gerstner, 2005; Fig. 4B). The presynaptic neurons have a firing rate of 10 Hz and refractory period of 1 ms. The STP parameters of the Tsodyks-Markram synapses were determined by fitting the induced postsynaptic responses to the data (Fig. 4C; see Methods). With this, the first-stage PFP facilitation of the model in the pre-pre, but not post-pre, ISI condition is predetermined (Fig. 4D, left).

**Figure 4:**
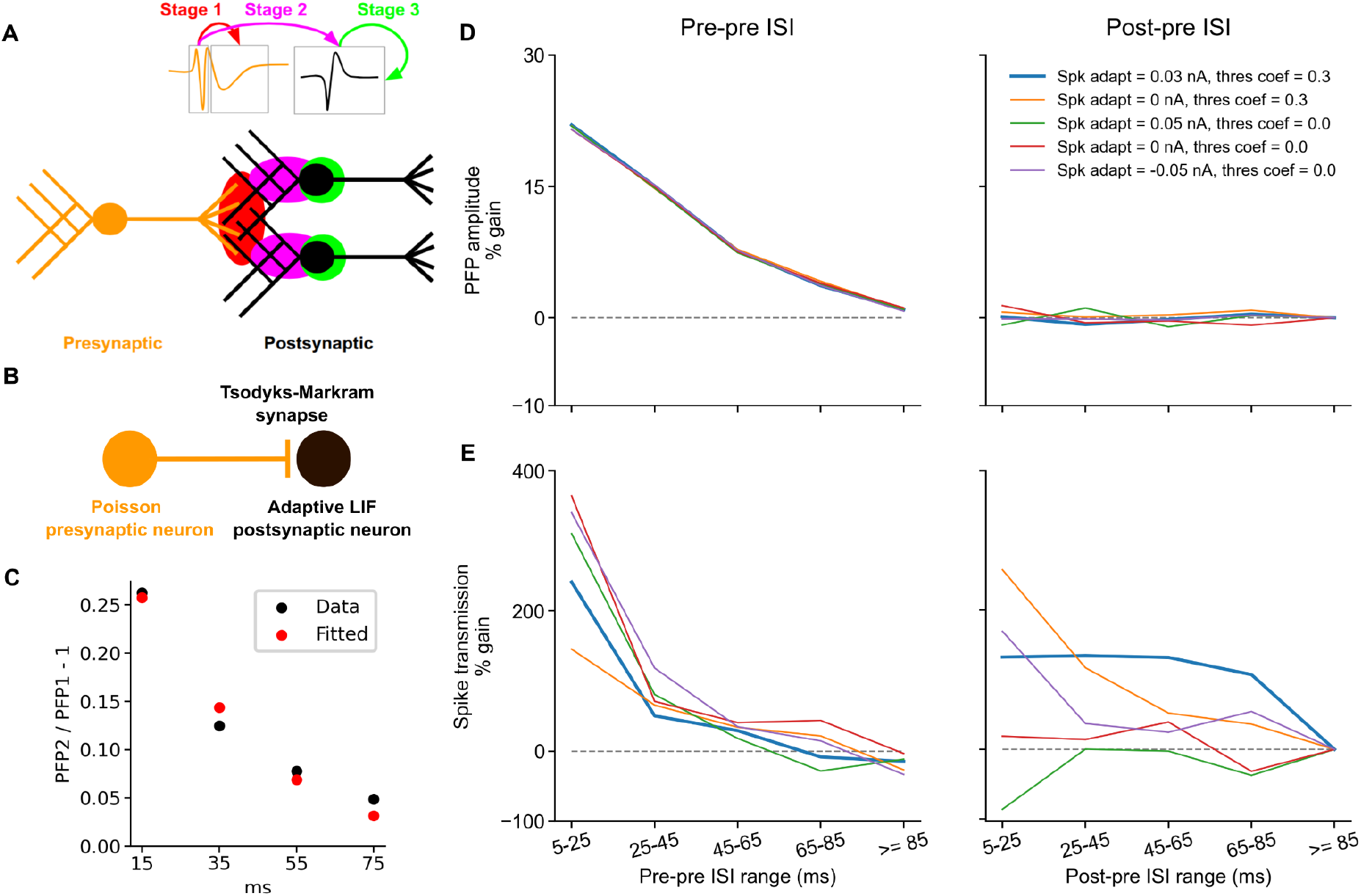
Three-stage short-term plasticity model. (A) A scheme illustrating the three-stage STP *in vivo*. The stage-1 STP (red) represents the dynamics of postsynaptic responses in response to presynaptic spikes, stage-2 STP (magenta) represents the dynamics of postsynaptic spiking in response to presynaptic spikes, and stage-3 STP (green) represents the dynamics of postsynaptic spiking in response to its recent spiking history. (B) A scheme illustrating the model setup with five Poisson presynaptic neurons and one adaptive LIF postsynaptic neuron, where each synapse implements the Tsodyks-Markram STP. (C) The synaptic STP of the model (red) was fitted to the median of data (black) in Fig. 1E. (D-E) The short-term facilitation of the model PFP (D) and spike transmission (E) for the pre-pre (left) and post-pre (right) ISIs. Different colors indicate models that use different mechanisms: basic LIF neuron (red), LIF neuron with dynamic threshold only (orange), LIF neuron with positive spike adaptation current only (green), LIF neuron with negative spike adaptation current only (purple), and LIF neuron with both dynamic threshold and positive spike adaptation current (blue). Each line indicates the median of the five presynaptic neurons. The basic LIF neuron (red) can already exhibit facilitation in the pre-pre ISI condition. Only the LIF neuron with both dynamic threshold and positive spike adaptation current (blue) can account for facilitation in both pre-pre and post-pre ISI conditions with the same trend as observed in the experiments.

Interestingly, given this predetermined pre-pre PFP facilitation, the second-stage characterizing spike transmission facilitation actually emerged naturally from the LIF model without any additional mechanism (Fig. 4E, left, red). This suggests that the basic LIF mechanism is already sufficient to account for the nonlinear enhancement of spike transmission, which may arise directly from PFP facilitation in the pre-pre ISI condition. That is, given the above model setup (Fig. 4B), the facilitations of the PFP and the pre-pre ISI (Figs. 1 and 2) could already be reproduced by the model (Figs. 4D-E, left, red). However, the longer-timescale post-pre spike transmission facilitation did not emerge from this model (Fig. 4E, right, red). This third-stage post-pre spike transmission facilitation emerged only after adding a dynamic threshold and positive spike adaptation current (Fig. 4E, right, blue; see Methods). The dynamic threshold with a positive threshold coefficient reduces the spiking threshold of the postsynaptic neuron by a fraction of the gap between the current threshold and the reset voltage every time the postsynaptic neuron emits a spike (see Methods), thereby reducing the required presynaptic input for the postsynaptic neuron to fire an action potential. In other words, the postsynaptic neuron becomes more excitable for a period of time after it fires.

The positive spike adaptation current in the adaptive LIF model has an effect for reducing the spiking of the postsynaptic neuron (Fig. 4E, right, green), whereas the negative spike adaptation current has the opposite effect of increasing the postsynaptic firing rate (Fig. 4E, right, purple). However, negative spike adaptation current alone without dynamic threshold (Fig. 4E, right, purple) failed to give rise to the observed trend of long-lasting post-pre facilitation of spike transmission (Fig. 3C). Similarly, an LIF model with only a dynamic threshold and no positive spike adaptation current also failed to give rise to the observed long-lasting post-pre facilitation trend from our experiments (Fig. 4E, right, orange). These two models both give rise to some degrees of post-pre spike transmission facilitation (Fig. 4E, right, purple and orange), but the facilitation curves do not match the observed experimental data (Fig. 3C). Given the exponential nature of these two mechanisms (see Methods), this suggests that there are putative opposing interactions between spike adaptation current and dynamic threshold that give rise to the semi-stable, longer-lasting post-pre short-term facilitation observed experimentally. In our model, we reproduced the experimental observations by using a small positive spike adaptation current of 0.03 nA with 20 ms time constant and threshold coefficient of 0.3 with 160 ms time constant.

## DISCUSSION

This study shows that there are three stages of short-term facilitation *in vivo* in the mouse retinocollicular pathway, where the spike transmission facilitates remarkably more than the synaptic input. The first and second stages are coupled to a large extent and have similar shorter-lasting dynamics in comparison to the third stage. Both the first and second short-term facilitation stages exhibit simple exponential decay, where the second stage exhibits a stronger facilitation compared to the first. Interestingly, the second stage emerges from the basic LIF neuron without any additional mechanism or parameter tuning (Fig. 4E, left, red), suggesting that the synaptic component plays a dominant role. On the contrary, the third stage is mainly associated with previous postsynaptic spiking activity, which is likely due to the nonsynaptic intrinsic excitability of the postsynaptic neuron as it is independent from the facilitation of synaptic inputs. Taken together, this implies that the synaptic and nonsynaptic components of the STP span over different timescales, where the synaptic component spans a shorter timescale than the nonsynaptic component. These facilitations reflect the coordination of synaptic and nonsynaptic components, revealing the information transfer dynamics from the retina to the SC. These results altogether provide an insight into STP *in vivo*, paving the way for finer understanding and predictions of circuit computation on a larger scale in the living brain.

### Simultaneous extracellular recording of PFP and spike transmission *in vivo*

Measuring STP *in vivo* by pairing single-cell measurements of both presynaptic and postsynaptic neurons has always been a challenging task. One of the key advantages of our *in vivo* tangential recording approach over *ex vivo* brain slice recordings is that STP can be measured more under physiological condition using visually-evoked responses instead of electrically-evoked responses. By using stimuli specific to the monitored pathway, it raises the confidence that the measured STP better mirrors the innate characteristics of the circuitry. Besides, the neurons in an intact brain are in a more natural environment than neurons in an acute slice. Although the variability of the presynaptic input strength could be high due to the ongoing STP induced by ongoing visual stimulus-driven activity, this variability should be safely canceled out by averaging across a large number of spikes throughout the recording.

In addition, with waveform classification, we can distinguish the responses of individual neurons and axons (Sibille et al., 2022a). This allows us to measure presynaptic neuronal spiking, STP of postsynaptic responses, and STP of spike transmission in the postsynaptic neuron simultaneously on a larger scale. With two additional controlled conditions, we showed that the PFP facilitation is unlikely to be biased by the postsynaptic spikes (Figs. S3 and S4). Another indirect evidence showing that the postsynaptic spikes are unlikely to contribute to the PFP is that there is no facilitation observed in the PFP of the post-pre ISI condition (Fig. 3D), even though there is a remarkable facilitation in the postsynaptic spiking activity (Fig. 3C). Since these synaptic and nonsynaptic components are generic to neurons in all pathways, it is conceivable that these STP stages likely exist in other brain regions, e.g., cortex, where this recording technique can be applied (Sibille et al., 2024).

### Facilitation in spike transmission probability is much larger than facilitation in PFP amplitude

Our results show that the facilitation of the spike transmission is much larger than the facilitation of the PFP (Figs. 1, 2, and S5). Even though the pre-pre ISI spike transmission gain was underestimated due to the use of average pre-pre ISI spike transmission, instead of the first RGC spike, as the baseline, the spike transmission facilitation is still around six times the PFP facilitation for the 5-25 ms pre-pre ISI range. In the context of long-term potentiation (LTP), the higher facilitation in the population spikes compared to the facilitation in the field EPSP (fEPSP) has been observed more than three decades ago (Bliss and Lømo, 1973; Taube and Schwartzkroin, 1988). Our results now suggest that this phenomenon already exists at the level of individual axons and individual postsynaptic neurons *in vivo*, where the PFP facilitation (Fig. 1E) is relatively small compared to the gain in the spike transmission (Fig. 2E). Notably, the facilitation of the spike transmission has a shorter decay time constant than the facilitation of the PFP (Fig. S5). Together, this suggests that, within a RGC-SC neuron pair, the gain in the postsynaptic spiking outputs evoked by the gain in postsynaptic responses is highly nonlinear, illustrated by the decreasing gradient of the correlations between percentage gain in the spike transmission and the PFP plasticity (Fig. 2F). In addition, the facilitation in spike transmission is more distinguished than the PFP facilitation in the convergent RGC spike pairs (Fig. S7), suggesting that this phenomenon is general to all concomitant inputs to the postsynaptic neuron, regardless of whether these inputs originate from the same presynaptic partner or from different ones.

### Input-independent facilitation in spike transmission

Spiking transmission has long been used as the proxy for inferring synaptic properties (Ahissar et al., 1992; Constantinidis and Goldman-Rakic, 2002; English et al., 2017; Henze et al., 2002). However, to our surprise, the postsynaptic spiking activity is facilitated with no obvious increase in the presynaptic input during the post-pre ISI condition (Fig. 3E), indicating that other mechanisms are at play. An explanation that can account for this is the dynamic threshold (Azouz and Gray, 2000), where the spiking threshold of the postsynaptic SC neurons might change based on the past activity of that postsynaptic neuron. Indeed, adding a dynamic threshold mechanism to the adaptive LIF model helps to reproduce the experimental observation (Fig. 4E, right, blue), which is not present in the basic LIF neuron (Fig. 4E, right, red) and adaptive LIF neuron without this mechanism (Fig. 4E, right, green). Furthermore, the LIF neuron with either negative spike adaptation current alone (Fig. 4E, right, purple) or with dynamic threshold alone (Fig. 4E, right, orange) could produce the stage-3 post-pre spike transmission facilitation but with a different dynamics than the one observed experimentally. Rather, the post-pre spike transmission facilitation that has an intriguing steady, longer-lasting dynamics (Fig. 3C) could potentially be explained by the interplay between spike adaptation current and dynamic threshold. Presumably, the more rapid decay of positive spike adaptation current induces a short decrease in excitability (Fig. 4E, right, green), which effectively cancels out the initial large facilitation due to the dynamic threshold, whereas the slower decaying dynamic threshold gives a long increase in excitability (Fig. 4E, right, orange). Together, dynamics threshold and positive spike adaptation current give rise to a long and steady post-pre facilitation of spike transmission independent of the synaptic input (Fig. 4E, right, blue), similar to the enhanced EPSP-spike coupling effect (Mahon et al., 2003; Tominaga and Tominaga, 2016). This synergistic effect between the faster dynamics of spike adaptation current and the slower dynamics of dynamic threshold give rise to the overall longer-lasting post-pre spike transmission facilitation, suggesting that there could be distinct STP effects for the synaptic and the spike transmissions, where spike transmission likely encompasses both synaptic and nonsynaptic components.

Such a phenomenon has been reported in the context of LTP, where Bliss and colleagues observed a potentiation in the population spike amplitude without noticeable increase in the fEPSP (Bliss and Gardner-Medwin, 1973; Bliss and Lømo, 1973). They also noticed a decrease in the population spike latency, suggesting that the population spike potentiation is either due to a decrease in the spiking threshold (Chavez-Noriega et al., 1990) or increase in tonic activity from other pathways. With the advantage of measuring the spiking activity of individual postsynaptic neurons, we show that the facilitation in the spike transmission in our study is highly dependent on the interval to the preceding postsynaptic spike, indicating that a decrease in spiking threshold in the postsynaptic neuron is more likely to happen in our case than the tonic activity from other pathways. This spike transmission facilitation in post-pre ISI condition is unlikely to be due to the inputs from other pathways, e.g. the corticotectal or the deeper-to-superficial SC connections, because those inputs need to be very precise temporally, strong, and have specific structure in order to contribute considerably to the time window used for detecting monosynaptic connections (Stevenson, 2023). However, we cannot rule out the possibility that the facilitation observed in the post-pre ISI condition could be due to the circuit-wide mechanisms, such as amplification within the SC (Shi et al., 2017), or brain states that modulate the gain of the SC neurons by changing the background synaptic inputs (Chance et al., 2002; Ferguson and Cardin, 2020).

This transient enhancement of postsynaptic firing without change in presynaptic input has probably a nonsynaptic origin. The nonsynaptic component is likely due to the change in the intrinsic excitability of the neuron, which is challenging to study in the context of STP. The intrinsic excitability can alter the input-output function of a neuron by different mechanisms, e.g., alteration of excitation-inhibition balance (Marder and Buonomano, 2003), modulation of the GABAergic system (Tominaga and Tominaga, 2016), and modification of resting membrane potential or shifting of membrane channel sensitivity that leads to the adaptation in spiking threshold (Debanne et al., 2019). As shown in our model, the reduction in spiking threshold is one potential mechanism for the post-pre facilitation, possibly induced by use-dependent inactivation of voltage-dependent, slowly inactivating A-type potassium current (Mahon et al., 2003). Another potential mechanism is the change of resting membrane potential, which has been shown to affect the subsequent firing rate (Sánchez-Aguilera et al., 2014). Both mechanisms could be at play, but it remains challenging to distinguish these two in the absence of pharmacological intervention. In the model, we did not simulate the repolarization, the resting membrane potential was fixed and the membrane potential was immediately reset to the resting membrane potential once the postsynaptic neuron spikes. Therefore, our model only considered the changing threshold. Given the same presynaptic inputs, the postsynaptic spiking in the adaptive LIF neuron depends on the gap between the resting membrane potential and the threshold, therefore changing either of them using the same time constant will produce equivalent effects.

### Higher input spike rates slightly enhance the spike transmission gain in pre-pre but not post-pre ISI conditions

The enhancement of raw spike transmission gain in the pre-pre ISI condition with high input spike rate (Fig. S8D) is likely because the membrane potential of the postsynaptic SC neuron is closer to the spiking threshold for higher input spike rates than lower input spike rates. However, the general trend of facilitation of different pre-pre ISI groups is robust across different input spike rates, where the facilitation in the 5-25 ms is much higher than other ISI ranges (Fig. S8C). This is likely because the paired-spike facilitation in the convergent inputs from different RGCs (Figs. S7C and H) is much smaller than the paired-spike facilitation of the same RGCs (Figs. 1D and 2E), therefore the paired-spike facilitation of the same RGCs still dominates in the presence of other convergent RGC inputs. The input-spike-rate independent effect in the post-pre ISI condition (Fig. S8F) suggests that the facilitation of the spike transmission in the post-pre ISI condition is likely not stemming from the presynaptic RGCs, consistent with the observation that the PFP remains unchanged in the post-pre ISI condition (Fig. 3D), but the exact mechanism is unclear. Further investigations are needed to uncover the underlying mechanisms.

### Limitations of the current study

Despite the fact that this approach allows us to simultaneously measure PFP and postsynaptic neuronal spiking extracellularly, this approach also has some limitations. Although we could measure the PFP induced by a single axon, the PFP is the combined response of multiple postsynaptic sites likely originating from dendrites of different postsynaptic neurons that are connected to the axon (Sauvé et al., 1995), and it is not yet known how to reliably disentangle the dendritic response of a single postsynaptic neuron. Also, the precise molecular mechanisms, both presynaptic and postsynaptic, that are responsible for the observed short-term facilitation cannot be determined without more thorough investigation using pharmacological applications. Lastly, we do not distinguish different RGC cell types (Baden et al., 2016) and SC cell types (Liu et al., 2023) in this study, and it would be an important aspect to further refine our understanding of the STP on the cell-type specific level in future studies.

## Conclusions

STP determines the dynamical evolution of single-neuron activity in relation to the presynaptic and postsynaptic partners; it is a key factor that governs how neural information is routed in the circuits. STP also plays a key role in neural coding by exerting a strong influence on both spike timing and rate. With the means to directly measure *in vivo* synaptic signals, we can decompose synaptic from nonsynaptic components, which enables the study of their relationships and thus the classification of different STP stages. This provides a comprehensive view on how information is gated in neural circuits at both the synaptic and neuronal levels, giving insights into how neurons and circuits compute. In short, the diverse STP stages *in vivo* revealed by this study offer a simple yet effective way for understanding the circuit dynamics in living brains, and potentially in behaving animals.

## METHODS

### Preparation, animals, surgery, and histology

#### Animals

All experiments were conducted following the guidelines of the local authority (Landesamt für Gesundheit und Soziales - LAGeSo Berlin - G0142/18). Maximal care was provided to reduce the animal usage and their discomfort during all experiments. Adult male mice (C57BL/6J, *n* = 24) were obtained from the local breeding facility (Charité-Forschungseinrichtung für Experimentelle Medizin) and Charles-River Germany.

#### Surgery

Induction was performed using 2.5% isoflurane in oxygen (Cp-Pharma G227L19A). Once stabilized, the animal was placed in a stereotactic frame (Narishige) with a closed-loop temperature controller (FHC-DC) for surgical procedure. During surgery, the eyes were protected with eye ointment (Vidisic) and the isoflurane level was slowly lowered (0.7-1.5%) while making sure that responses to tactile stimulation and vibrissa twitching remain absent. A craniotomy was carried out shortly after constructing the dental cement-based crown (Paladur, Kuzler) used for fixing the head post and grounding. Once surgery and craniotomy were finished, the animal was transferred onto the recording table with a heating pad in place to keep the animal warm for the rest of the experiment. The experiment using visual stimuli (described below) was recorded once the Neuropixels probe is inserted tangentially and stabilized within the mouse SC.

#### Histology

Once the recording finished, the probe was removed, coated with DiI (Abcam-ab145311 diluted in ethanol), and re-inserted in the same tissue location for 5 minutes. The animal was then sacrificed with excess isoflurane (>4%). Phosphate buffer saline solution (PBS) was used for cardiac perfusions, followed by 4% paraformaldehyde (PFA) in PBS. The brains were kept in 4% PFA at least overnight and stored in PBS until histological slicing (Leica VT1200 S). DAPI-Fluoromount-G (70-100 µm slices, Biozol Cat. 0100-20) was used to mount the brain slices.

#### 3D location reconstruction

The Neuropixels probe track was reconstructed in 3D using SHARP-track (Shamash et al., 2018) and the Allen Mouse Brain Common Coordinate Framework (Steinmetz et al., 2019). The SC visually driven responses were used as landmarks to align and scale the estimated recording site positions.

### Visual stimulation

#### Visual stimuli

All visual stimuli exposed to the animals were generated in Python using PsychoPy toolbox (Peirce, 2008). A transistor-transistor logic (TTL) signal was generated and time-locked to the screen update for every stimulus onset. To cover a large part of the visual field, the visual stimuli were projected on a spherical dome (projector (NEC ME331W, refresh rate = 60 Hz, mean luminance = 110 cd/m^2^, Gamma corrected, *n* = 20 mice) via a plexiglass reflecting half bowl (Modulor, 0260248). The inner part of the dome was covered with a layer of broad-spectrum reflecting paint (Twilight-labs) to increase the brightness of the reflected image (Denman et al., 2017). The projected image was warped using the meshmapper software (http://paulbourke.net/dome/meshmapper) to compensate for the deformation of the image reflections. In a subset of experiments for pharmacological controls (*n* = 4 mice), an LCD display (Dell S2716DG, refresh rate = 120 Hz, mean luminance = 120 cd/m^2^, Gamma corrected) was used instead of the spherical dome. The LCD display was positioned on the side of the animal and aligned to the pupil resting position in order to maximize the number of visually driven channels.

### Electrophysiological recordings

#### Neuropixels recording

The neurons in the mouse SC were recorded with Neuropixels probes (Phase 3a and Phase 3B1) (Jun et al., 2017) using the Open Ephys software (www.open-ephys.org). All recordings were done with the PXIe system (National Instrument NI-PXIe-1071), which stores the extracellular signals in the local field potential (0.5-500 Hz) and the action potential bands (0.3-10 kHz).

#### SC tangential insertion

The Neuropixels probe was inserted using a micromanipulator system (NewScale, MPM-M3-LS3.4-15-XYZ Upright). Stereotactic coordinates were described in reference to lambda in the medio-lateral (ML), dorso-ventral (DV), and antero-posterior (AP) axes. All angles were defined in reference to the azimuthal plane at lambda (Paxinos and Franklin, Nixdorf 2007 stereotaxic atlas). The Neuropixels probe was inserted tangentially (15 to 25 deg) into the SC from the back (500 to 1200 μm ML, -100 to -500 μm DV, -100 to -300 μm AP from lambda) and remained parallel to the sagittal plane during the experiment. The Neuropixels probe was gradually driven >4 mm into the target tissue followed by a small withdrawal of 20 to 50 µm to release accumulated mechanical pressure. The probe was then left for ∼10-20 minutes to settle. To ensure that the probe is located within the area that is being visually stimulated, the receptive fields of the multi-unit-activity were mapped using a customized script (Sibille et al., 2022b) before any recording.

#### Waveform extraction and spike sorting

The extracellular neuronal signals were spike-sorted using Kilosort (2 and 2.5; Pachitariu et al., 2016). Manual curation using Phy2 (https://github.com/cortex-lab/phy) was performed on the Kilosort output. To avoid biases from noisy outputs, quality metrics thresholding (isolation distance > 10, refractory period violation of ±1.5 ms for <0.01% of all spikes), removal of unstable clusters, and controls for double-counted spikes were performed after the manual curation and before any further analysis. The identification of RGC axonal action potential waveforms was based on the characteristic presence of double negative peaks within 3 ms, where all RGCs were detected by Kilosort based on the first negative peak of the waveform, corresponding to the extracellular potential evoked by the RGC axonal arbors in the SC (Sibille et al., 2022a). This ensured the detection of RGC spikes independent of any postsynaptic plastic effect.

### Interspike interval (ISI) ranges

#### Pre-pre ISI

The pre-pre ISI is the ISI of two consecutive spikes of an RGC. For each RGC, if a pair of spikes has a dead time of at least 85 ms prior to the first spike, they will be categorized into one of the five ISI ranges: 5–25, 25–45, 45–65, 65–85, and ≥85 ms.

#### Post-pre ISI

The post-pre ISI of an RGC-SC neuron pair is the interval from the last occurring postsynaptic spike to the current presynaptic spike. To remove the potential effects of the pre-pre ISI, the post-pre ISI is valid only if there is a dead time of at least 85 ms before the presynaptic RGC spike time. The valid post-pre ISI was then categorized into one of the five ISI ranges: 5–25, 25–45, 45–65, 65–85, and ≥85 ms.

### Analysis of postsynaptic field potential

#### PFP based on ISI range

The postsynaptic field potential (PFP) is the extracellular postsynaptic potential originated from the activation of multiple postsynaptic sites evoked by the spike of their same presynaptic RGC partner (Sauvé et al., 1995; Sibille et al., 2022a). Its characteristic feature is a long lasting second negative deflection following the RGC axonal spike, whose postsynaptic nature has been confirmed pharmacologically using a cocktail of synaptic blockers (Sibille et al., 2022a). The one-dimensional waveform of the PFP of an RGC was obtained by averaging 11 channels of the recalculated multi-channel waveform centered at the best channel defined by the Kilosort’s output (Fig. 1C). The PFP for each pre-pre ISI range was obtained from averaging the RGC spike waveforms within the same pre-pre ISI range. The PFPs evoked by the first and second spikes of the RGC spike pairs (with a valid dead time) within a pre-pre ISI range were averaged separately. For the condition without tailgating postsynaptic SC spike, the waveforms of the first and second RGC spikes of each pre-pre ISI range were recalculated by removing RGC spikes that have any recorded SC spike occurred within the [-3, 6] ms window around the RGC spikes. Similar to the pre-pre ISI condition, the PFP for each post-pre ISI range was obtained from the average RGC spike waveform of the post-pre ISI range.

#### PFP plasticity

Three variables were used for quantifying the PFP plasticity, namely PFP amplitude, peak-to-peak PFP amplitude, and PFP slope (Fig. S3A). The amplitude of the PFP was defined as the difference from the RGC axonal spike waveform baseline to the minimum point of the PFP (Fig. 1D). The peak-to-peak PFP amplitude was defined as the amplitude from the previous peak to the minimum point of the PFP and the PFP slope was defined as the slope from 0.2 to 0.8 fraction of the peak-to-peak PFP (Fig. S3A). The PFP of an RGC was included if (i) the number of RGC spike pairs for every ISI range is at least 500 (50 for the spike triplets and quadruplets in Fig. S2B, 200 for the muscimol group in Fig. S4), (ii) its amplitude evoked by the second RGC spike at ≥ 85 ms pre-pre ISI is at least 5 μV in size, and (iii) the ratio of the PFP amplitude to the peak-to-peak PFP size is at least 0.3.

The plasticity in the PFP of a pre-pre ISI range, *T*, was defined as the percentage gain of PFP amplitude evoked by the second RGC spike, *A*^2*nd,T*^, over the PFP amplitude evoked by the first RGC spike, 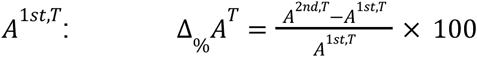 *T* ∈ {[5, 25), [25, 45), [45, 65), [65, 85), [85, ∞)} ms. For convergent inputs from different RGCs (Fig. S7), the first and second PFP amplitudes are first normalized to their corresponding PFP amplitude at ≥ 85 ms before computing the percentage: 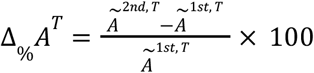, where 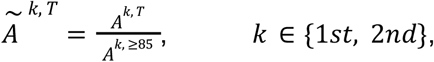, and *T* ∈ {[5, 25), [25, 45), [45, 65), [65, 85)} ms. For the post-pre ISI condition, the change in the PFP amplitude of an ISI range was defined as the percentage of PFP amplitude of a test ISI range, *A*^*T*^, over the PFP amplitude of the baseline ISI range at ≥ 85 ms, *A*^≥85^ : 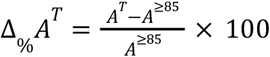, *T* ∈ {[5, 25), [25, 45), [45, 65), [65, 85)} ms.

### Cross-correlation analyses

#### Baseline corrected cross-correlation

Spike transmission was estimated from the spike train cross-correlogram (CCG) of an RGC-SC neuron pair. For unbiased estimation of the spike transmission, a slow co-modulated baseline, λ(*m*), was computed for each CCG, *C*(*m*), and subtracted from the CCG to get the baseline-corrected CCG, *Z*(*m*) = *C*(*m*) − λ(*m*) (English et al., 2017; Stark and Abeles, 2009). This slowly modulated baseline was computed by convolving a partially hollow Gaussian kernel (hollow fraction = 0.6, standard deviation = 10 ms, kernel length = 15 ms, bin width = 0.1 ms) with the CCG. The section at both ends of the CCG, with length equals to half the kernel length (7.5 ms), were duplicated, flipped, and symmetrically appended to the CCG when computing the CCG baseline to prevent any edge effects (Stark and Abeles, 2009). For the CCG of the post-pre ISI condition, there is a gap of the size of the smallest ISI, *T*_*min*_, within the post-pre ISI range. For example, there is a gap of 5 ms for the post-pre ISI range of 5-25 ms. This gap, with time window (− *T*_*min*_, 0] ms, was filled with the duplicated and flipped values from the post-pre ISI CCG window of [5, 5 + *T*_*min*_ ) ms when computing the post-pre ISI CCG baseline.

#### Connection detection

Connections are estimated from the excess count in the *m*th time lag of the CCG in comparison to the baseline using Poisson distribution with a continuity correction (English et al., 2017; Stark and Abeles, 2009). The RGC-SC pairs were labeled as connection if the probability, *P*, of obtaining an observed (or higher) count *n* in the *m*th time lag of the CCG, given the λ(*m*), is less than 0.001 for at least eight consecutive lag bins (bin width = 0.1 ms) within the lags of 0.8 - 2.8 ms.

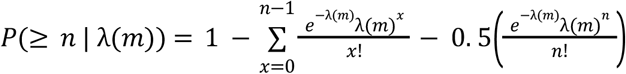

For analyses of the pre-pre and post-pre ISI conditions, the average CCG across ISI ranges was considered as having a significant effect if *P* < 0.01 for at least five consecutive lag bins. For the pre-pre ISI condition, the average CCG was computed from only the second RGC spikes of all ISI ranges.

### Spike transmission probability

#### Efficacy of synaptic connection

The spike transmission probability, *P*_*spike*_, was computed from the rectified baseline-corrected CCG, *Z*^+^ (*m*), averaged over *n* presynaptic spikes (English et al., 2017)

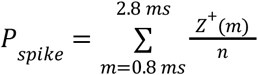

where .*Z*^+^ (*m*) = *max*(*Z*(*m*), 0)

H3>Plasticity of spike transmission:

The gain in spike transmission probability, *G*, was defined as the excessive spike transmission probability within lag window of 0. 8 − 2. 8 ms in a test group in comparison to the baseline group: 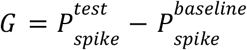 . The percentage gain in spike transmission probability, *G*_%_, was defined as the percentage gain in spike transmission probability relative to the average spike transmission probability for the respective condition, 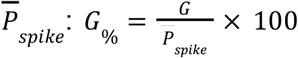 . Average spike transmission probability was used as the baseline instead of transmission probability of the first spike because the transmission probability of the first spike of some RGC-SC pairs was very small, which might result in an erroneous calculation of gains. The RGC-SC pair was included if (i) the PFP of the RGC fulfilled the PFP inclusion criteria above, (ii) the overall connection efficacy from all RGC spikes is at least 0.01, and (iii) there is a significant effect in the average CCG across ISI ranges for the corresponding ISI condition described above.

For the pre-pre ISI condition, the gain in spike transmission probability was defined as the excessive spike transmission probability in the second spike compared to the first spike of the RGC spike pair within an ISI range: 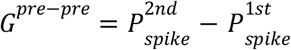. The percentage gain in *P*_*spike*_ for the pre-pre ISI condition, 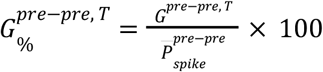, was defined as the percentage gain of an ISI range *T* ∈ {[5, 25), [25, 45), [45, 65), [65, 85), [85, ∞)} ms, *G*^*pre*−*pre, T*^, relative to the average spike transmission probability of the first and second spikes of all ISI ranges, 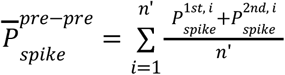, where *n*’ is the total number of valid spike pairs of an RGC. For convergent inputs from different RGCs of different synaptic strengths (Fig. S7), the two RGC spike trains were first combined, and then the percentage difference in spike transmission probability gain, 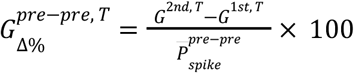, was computed instead, where 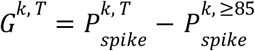, k ∈ {1*st*, 2*nd*}, and *T* ∈ {[5, 25), [25, 45), [45, 65), [65, 85)} ms is the convergent ISI range with the first spike of the ISI from one RGC and the second spike from another RGC.

For the post-pre ISI condition, gain in *P*_*spike*_ was defined as the excessive spike transmission probability in an ISI range, *T*, compared to the baseline ISI range at ≥ 85 ms: 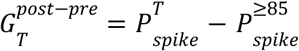, *T* ∈ {[5, 25), [25, 45), [45, 65), [65, 85)} ms. The percentage gain in *P*_*spike*_ for the post-pre ISI condition, 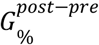, was defined as the percentage of 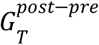 to the average spike transmission probability of all valid post-pre ISIs of an RGC-SC neuron pair.

### *Ex vivo* experiments and data analysis

#### Animal

Experiments were carried out according to the guidelines of the European Community Council Directives of 1 January 2013 (2010/63/EU). Mice (Mus musculus) were group housed on a 12-h light/dark cycle and all efforts were made to minimize the number of used animals and their suffering.

#### Slice preparation

Acute sagittal SC slices (400 µm) were prepared from 4-6 weeks-old C57BL/6 mice. Briefly, slices were cut at low speed (0.04 mm/s) and at a vibration frequency of 70 Hz in ice-cold oxygenated ACSF supplemented with sucrose (in mM: 87 NaCl, 2.5 KCl, 2.5 CaCl2, 7 MgCl, 1 NaH2PO4, 25 NaHCO3, 75 sucrose and 10 glucose, saturated with 95% O2 and 5% CO2). Slices were maintained in a storage chamber containing standard ACSF (in mM: 119 NaCl, 2.5 KCl, 2.5 CaCl2, 1.3 MgSO4, 1 NaH2PO4, 26.2 NaHCO3, and 11 glucose, saturated with 95% O2 and 5% CO2) at 34°C for at least 1 hour before recording.

#### Electrophysiological recordings

Slices were transferred in a submerged recording chamber mounted on an Olympus BX51WI microscope equipped for infrared differential interference microscopy and were perfused with standard ACSF at a rate of 2 ml/min at 34°C. Extracellular field potential (fEPSP) recordings were performed in SC superficial layers using glass pipettes (2-5 MΩ) filled with ACSF. Basal evoked postsynaptic responses were induced by stimulating SC afferents at 0.1 Hz. PPF was performed by delivery of two low stimuli (<10 mA) at various inter-pulse intervals (5-25, 30-50 and 60-100 ms). Repetitive stimulation was performed by delivery 2, 3 or 4 stimuli at 10 ms interval. Field potential recordings were acquired with Axopatch-1D amplifiers (Molecular Devices), digitized at 10 kHz, filtered at 2 kHz, and stored and analyzed on computer using pCLAMP 9 and Clampfit 10 software (Molecular Devices).

#### Statistics

Data are expressed as means ± SEM, unless otherwise stated. Repeated measures one-way ANOVA with Dunnett’s multiple comparison test was performed for PPF and repetitive stimulation comparisons.

### Short-term plasticity model

#### Model setup

The model consists of five Poisson presynaptic neurons and one leaky-integrate-and-fire (LIF) postsynaptic neuron with adaptation current (Brette and Gerstner, 2005) and dynamic threshold (Horn and Opher, 2000). The model was simulated for 1000 seconds with a time step of 0.5 ms. Each presynaptic neuron has a firing rate of 10 Hz with a refractory period of 1 ms.

#### Model synapse

The synapses implement Tsodyks-Markram model (Mongillo et al., 2008; Tsodyks and Markram, 1997), governed by the following kinetic equations for recovered (*B*(*t*)) and effective (*E*(*t*)) resources:

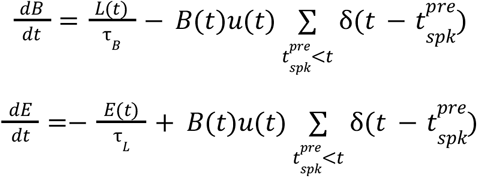

Where *L*(*t*) = 1 − *B*(*t*) − *E*(*t*) represents the inactive resources, τ_*B*_ = 28. 309 ms is the recovery time constant, τ = 1 ms is the inactivation time constant, δ(*t*) is the Dirac delta function, and 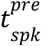 is the presynaptic spike times. The resource utilization dynamics is defined by

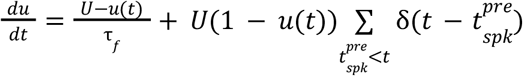

Where *U* = 0. 174 is the baseline utilization and τ_*f*_ = 25. 854 ms is the facilitating time constant. The Tsodyks-Markram parameters above were determined by fitting the median of data in Fig. 1E (Fig. 4C). The excitatory postsynaptic current (EPSC) induced by the *i*-th presynaptic neuron was then computed as *I*_*syn, i*_(*t*) = *A*_*SE*_*E*_*i*_(*t*), where *A*_*SE*_ = 3400 pA is the absolute synaptic efficacy that gives the maximal EPSC of a presynaptic neuron if all its resources are in the effective state. This EPSC was then used as the synaptic input current to the postsynaptic neuron with a synaptic delay of 1 ms.

#### Postsynaptic neuron

The membrane potential *V*(*t*) at time *t* of the postsynaptic neuron follows the dynamics:

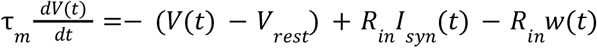

where τ_*m*_ = 10 ms is the membrane time constant, *V*_*rest*_ =− 65 mV is the resting potential, *R*_*in*_ = 150 MΩ is the input resistance, 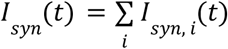 is the total synaptic input current of all presynaptic neurons at time *t*, and *w*(*t*) is the adaptation current. The dynamics of the adaptation current is as follows:

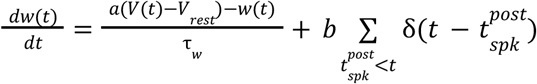

where τ _*w*_ = 20 ms is the adaptation time constant, *a* = 0 nS is the subthreshold adaptation, *b* = 0. 03 nA is the spike-triggered adaptation current, and 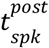 is the spike times of the postsynaptic neuron. Since *a* = 0 nS, the adaptation dynamics is reduced to:

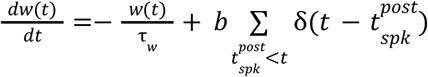

The membrane potential is reset to *V*_*r*_ = *V*_*rest*_ =− 65 mV once the membrane potential reaches the threshold. We implemented a variant of dynamic threshold (Horn and Opher, 2000) governed by the following dynamics:

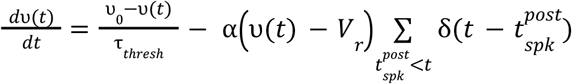

where τ_*thresh*_ = 160 ms is the dynamic threshold time constant, *υ*_0_ =− 45 mV is the resting threshold, and α = 0. 3 is the dynamic threshold coefficient.

#### Model analyses

The PFP amplitudes were estimated from the EPSP trace of each presynaptic neuron. The spike train cross-correlation (Fig. S9) was computed as described above but with a bin width of 0.5 ms, which is the model time step. All other analyses were performed using the same procedures as the data analyses described above.

## ACKNOWLEDGEMENTS

We thank Carolin Gehr and Tatiana Lupashina for helpful discussions during the project. We thank Francois David for comments on the manuscript. This work was supported by DFG grants KR 4062/4-1, KR 4062/4-2, KR 4062/5-1, and KR 4062/6-1 (J.K.) and the Major Research Program of PSL Research University “PSL-Neuro” launched by PSL Research University and implemented by ANR (ANR-10-IDEX-0001) (N.R.).

## AUTHOR CONTRIBUTIONS

K.-L.T., J.S., and J.K. designed the study; J.S. designed and conducted *in vivo* experiments and curated the data; K.-L.T. analyzed the *in vivo* data; E.D. and N.R. designed and conducted the *ex vivo* experiments; E.D. analyzed the *ex vivo* data; K.-L.T. implemented and studied the computational model; and K.-L.T. wrote the manuscript with inputs from all authors.

## SUPPLEMENTARY MATERIALS

**Figure S1:**
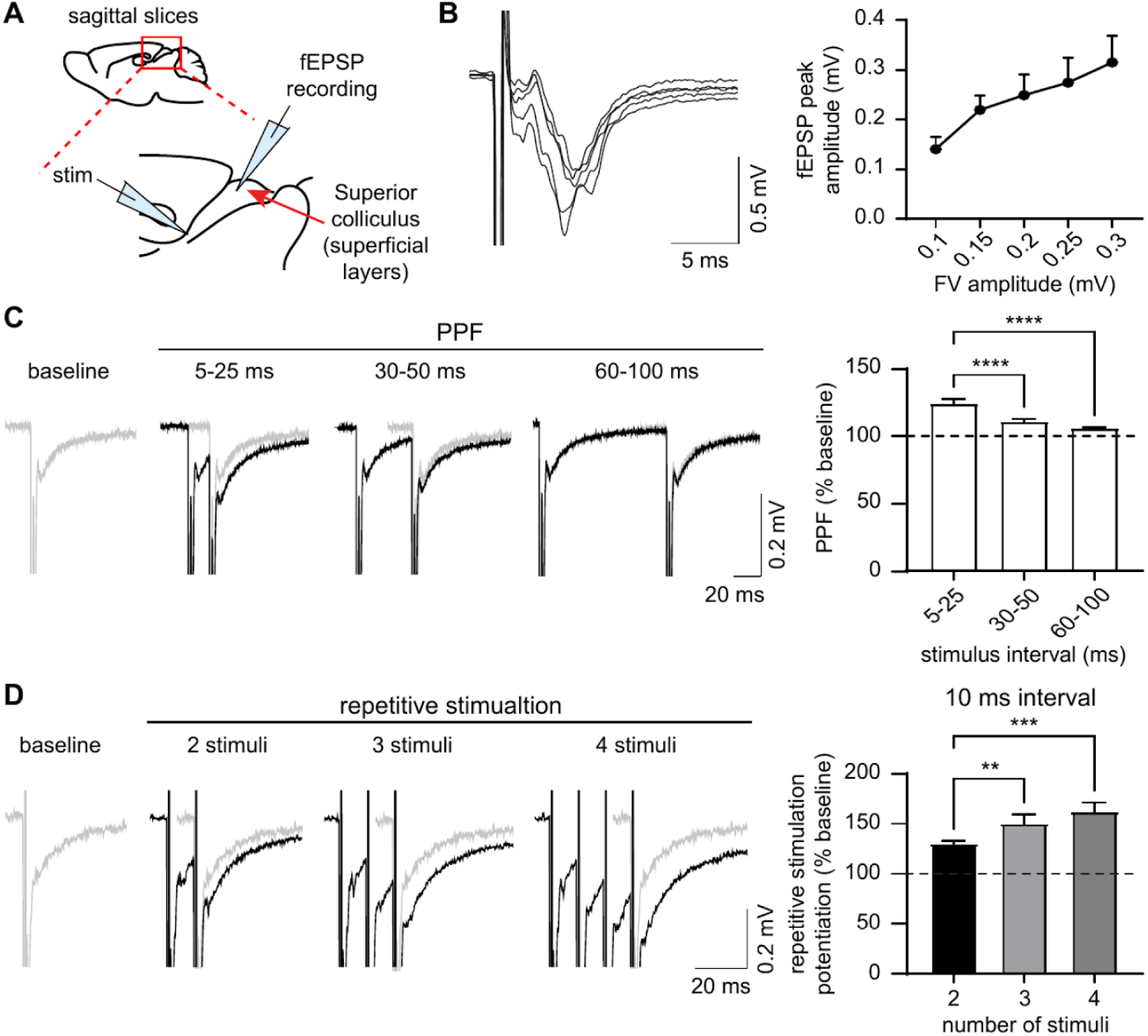
*Ex vivo* SC slices exhibit short-term facilitation in paired-pulse and repetitive stimulation protocols. (A) Schematic representation of electrode positions used in brain sagittal slices to record fEPSPs in the superficial layers of the SC evoked by electrical stimulation of SC afferents. (B) Input-output curves for basal synaptic transmission. Left: representative recordings in wild-type mice. Scale bars, 5 ms, 0.5 mV. Right: quantification of the fEPSP peak amplitude for different fiber volley amplitudes after stimulation (*n* = 7 slices from 3 mice). The data are represented as the mean ± sem. (C) Left, representative traces (black) of paired-pulse facilitation (PPF) with different inter-pulse intervals (5-25, 30-50 and 60-100 ms) recorded in the superficial layers of the SC after electrical stimulation of SC afferents. The baseline response to 0.1 Hz stimulation is shown in grey. Scale bars: 20 ms, 0.2 mV. Right, quantification of the PPF as % of the baseline response (*n* = 7 slices from 3 mice; *p* < 0.0001, RM-one way ANOVA). (D) Left, representative traces (black) of the response to repetitive stimulation (2, 3 and 4 stimuli at 10 ms interval) recorded in the superficial layers of the SC after electrical stimulation of SC afferents. The baseline response to 0.1 Hz stimulation is shown in grey. Scale bars: 20 ms, 0.2 mV. Right, quantification of the repetitive stimulation-induced potentiation as % of the baseline response (*n* = 6 slices from 3 mice; *p* = 0.007, RM-one way ANOVA). Asterisks indicate statistical significance (**, *p* < 0.01; ***, *p* < 0.001; ****, *p* < 0.0001).

**Figure S2:**
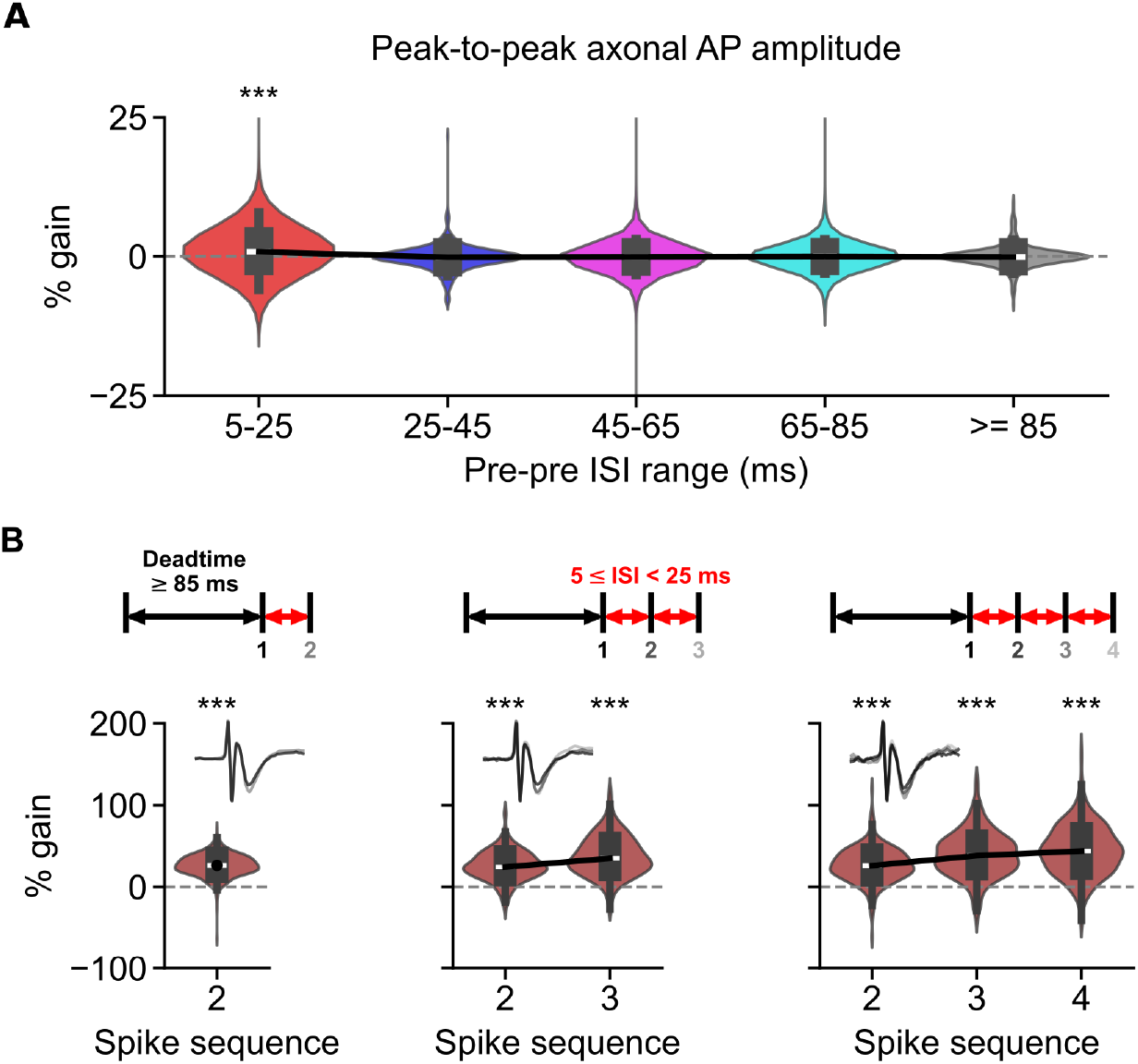
Axonal AP amplitude is not affected by the pre-pre ISI whereas consecutive RGC spikes induce lasting PFP facilitation. (A) The distributions of peak-to-peak axonal AP amplitude for different ISI ranges, where the peak-to-peak axonal AP amplitudes are similar across different ISI ranges. The black line indicates the median, the box inside the violin plot represents the Q1 to Q3 of the data (Q1: -0.75%, -0.99%, -0.85%, -0.75%, -0.8%; median: 0.85%, -0.12%, -0.05%, 0.03%, -0.07%; Q3: 2.68%, 0.7%, 0.72%, 0.76%, 0.71%; *p* = 1.91 x 10^-11^, 0.059, 0.199, 0.667, 0.293 for ISI ranges of 5-25, 25-45, 45-65, 65-85, and ≥85 ms, two-sided Wilcoxon signed-rank test, *n* = 498 RGCs, *n* = 27 experiments, *n* = 24 mice). Although the change of the RGC axonal AP amplitude of the 5-25 ms ISI range is significant compared to the baseline, its effect size is very small (Cohen’s d = 0.00909) and thus negligible. (B) Two (same as Fig. 1E), three (Q1: 14.68%, 21.21%; median: 24.4%, 35.04%; Q3: 37.06%, 53.58%; *p* = 2.41 x 10^-71^, 3.61 x 10^-76^ for the second and third spikes, two-sided Wilcoxon signed-rank test, *n* = 484 RGCs, *n* = 27 experiments, *n* = 24 mice), and four (Q1: 14.49%, 21.93%, 22.51%; median: 26.11%, 38.31%, 43.96%; Q3: 39.53%, 56.62%, 65.1%; *p* = 2.28 x 10^-49^, 8.72 x 10^-51^, 1.27 x 10^-52^ for the second, third, and fourth spikes, two-sided Wilcoxon signed-rank test, *n* = 343 RGCs, *n* = 25 experiments, *n* = 22 mice) consecutive RGC spikes within ISI of 5-25 ms induced lasting PFP facilitation. Top: scheme showing the different number of incoming presynaptic spikes after a dead time of at least 85 ms. Bottom: PFP amplitudes normalized to the PFP of the first spike. The black line indicates the median, the box inside the violin plot represents the Q1 to Q3 of the data.

**Figure S3:**
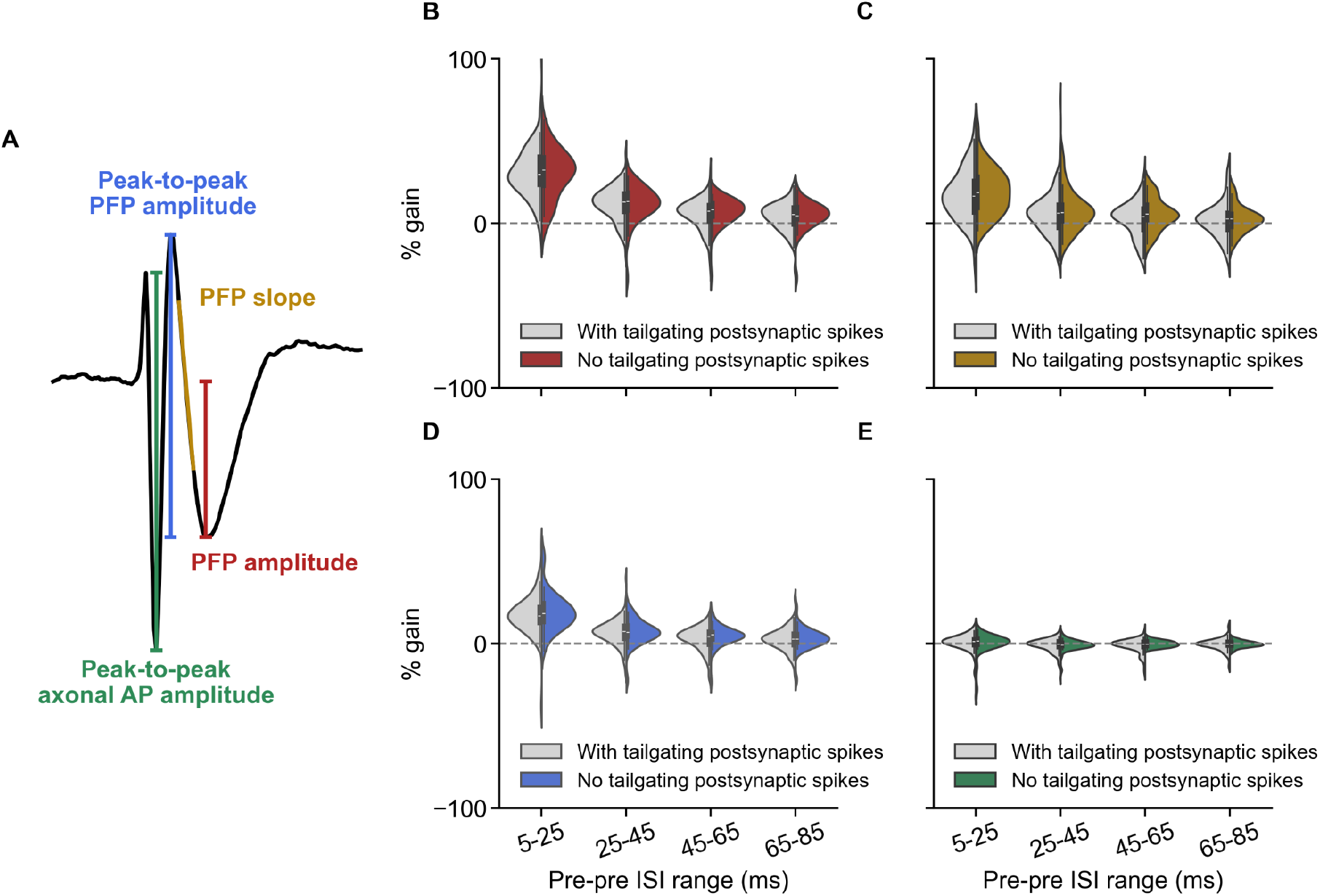
No significant difference in the PFP facilitation with and without tailgating postsynaptic SC spikes at the lag window [-3, 6] ms around the RGC spike time. (A) Scheme showing different variables of the 1-D RGC waveform being quantified. (B-E) The percentage change of different RGC waveform variables of the second RGC spike in different pre-pre ISI ranges in comparison to the baseline ISI range of ≥ 85 ms. No obvious difference observed between the waveforms with (white) and without (colored) tailgating postsynaptic SC spikes for facilitation in PFP amplitude (B; *p* = 0.904, 0.954, 0.365, 0.617), PFP slope (C; *p* = 0.363, 0.523, 0.251, 0.718), peak-to-peak PFP amplitude (D; *p* = 0.483, 0.872, 0.282, 0.602), and peak-to-peak axonal AP amplitude (E; *p* = 0.922, 0.668, 0.401, 0.738). For all variables, a two-sided Wilcoxon rank-sum test was used. For the group with tailgating postsynaptic SC spikes: *n* = 110 RGCs, *n* = 10 experiments, *n* = 9 mice. For the group without tailgating postsynaptic SC spikes: *n* = 37 RGCs, *n* = 5 experiments, *n* = 5 mice.

**Figure S4:**
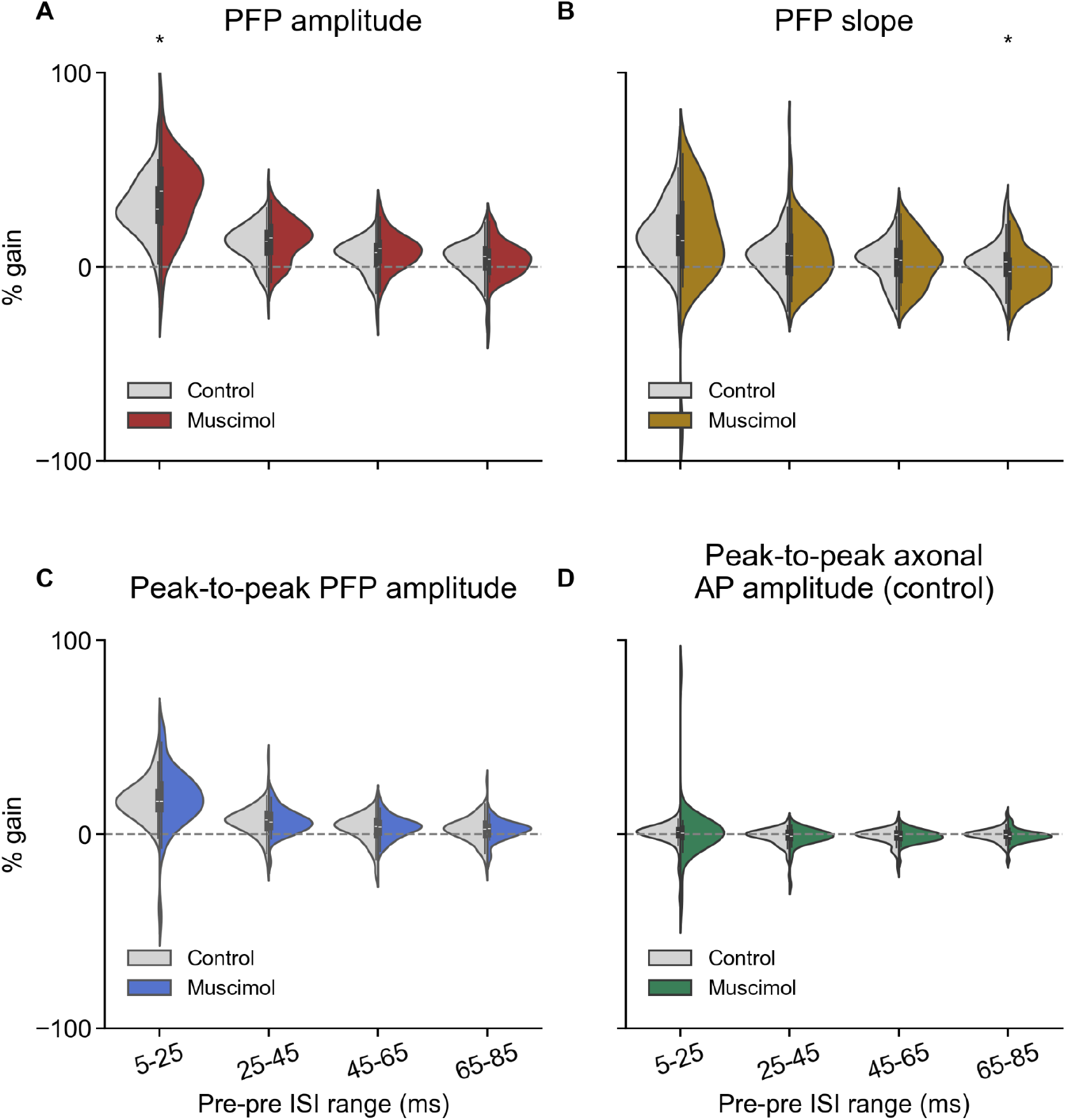
No obvious difference in the PFP facilitation between the control and with muscimol application in the SC. (A-D) The percentage change of different RGC waveform variables of the second RGC spike in different pre-pre ISI ranges in comparison to the baseline ISI range of ≥ 85 ms. No obvious difference observed between the controlled waveforms (white) and waveforms under muscimol condition (colored, 2.5 mM) for facilitation in PFP amplitude (A; *p* = 0.046, 0.258, 0.23, 0.401), PFP slope (B; *p* = 0.914, 0.901, 0.974, 0.016), peak-to-peak PFP amplitude (C; *p* = 0.541, 0.504, 0.974, 0.675), and peak-to-peak axonal AP amplitude (D; *p* = 0.614, 0.507, 0.598, 0.101). For all variables, a two-sided Wilcoxon rank-sum test was used. For the control group: *n* = 110 RGCs, *n* = 10 experiments, *n* = 9 mice. For the muscimol group: *n* = 44 RGCs, *n* = 3 experiments, *n* = 3 mice.

**Figure S5:**
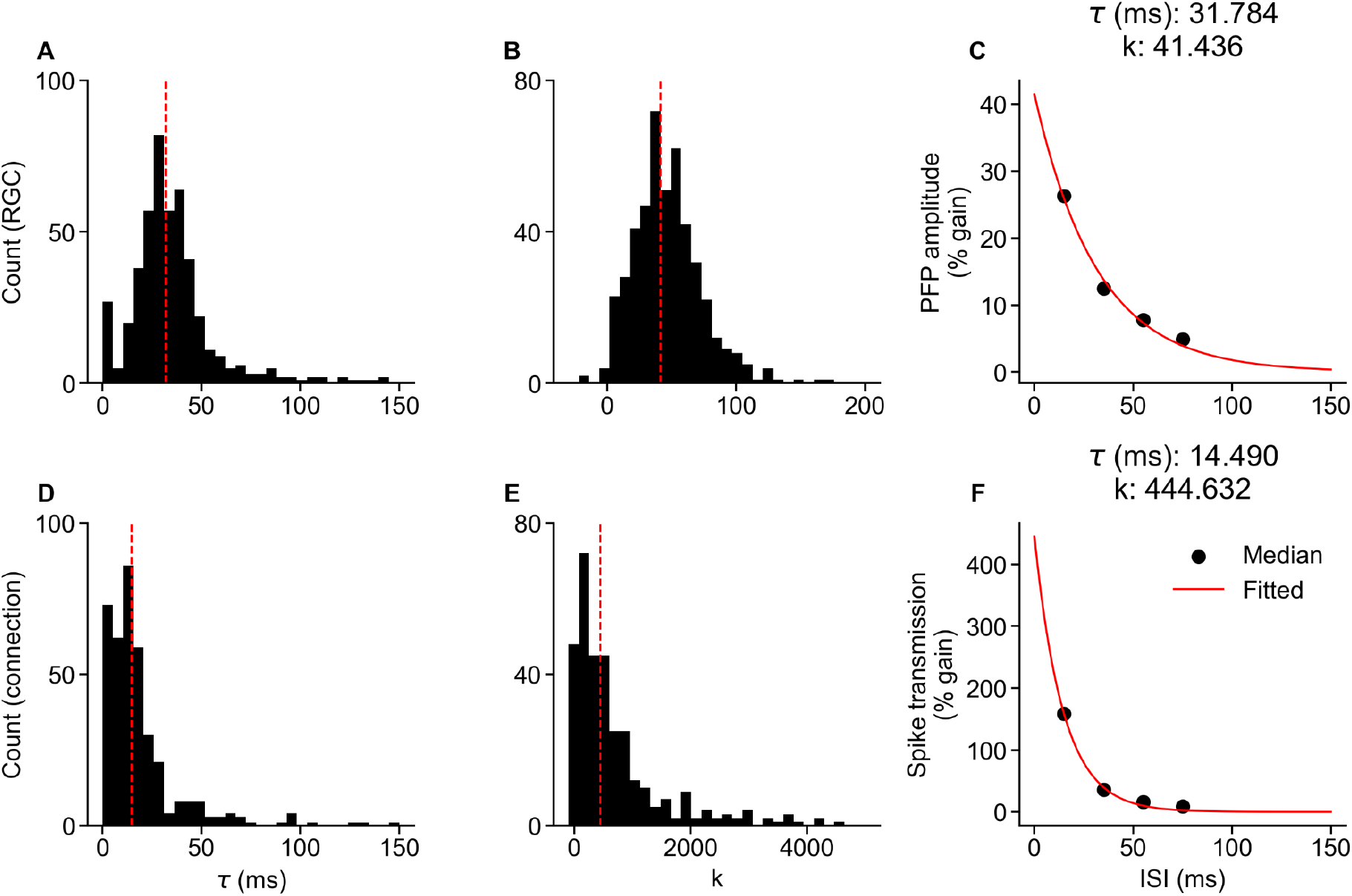
Short-term facilitation of the PFP is smaller but longer-lasting compared to the spike transmission. The STP of the PFP amplitude and spike transmission were fitted to an exponential decay function of the form 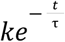, where *k* is the initial value of the exponential decay function at *t* = 0 ms, *t* ∈ {15, 35, 55, 75} ms is the midpoint of each pre-pre ISI group (not including the baseline group of ≥85 ms), and τ is the decay time constant of the STP effect. Negative *k* indicates depressing effect whereas positive *k* indicates facilitating effect. (A, D) The distribution of decay time constant τ of the STP for the percentage gain in PFP amplitude (A; *n* = 498 RGCs, *n* = 27 experiments, *n* = 24 mice) and the percentage gain in spike transmission (D; *n* = 406 pairs, *n* = 224 RGCs, *n* = 21 experiments, *n* = 20 mice). The dashed vertical red line is the τ of the median data. (B, E) The distribution of *k* for the PFP amplitude percentage gain (B) and the spike transmission percentage gain (E). The dashed vertical red line is the *k* of the median data. (C, F) The median data (black dots) and the fitted exponential decay function (red line) for the PFP amplitude percentage gain (C) and the spike transmission percentage gain (F). The τ and *k* of the fitted exponential decay function are corresponding to the vertical dashed line in A-B (PFP amplitude percentage gain) and D-E (spike transmission percentage gain).

**Figure S6:**
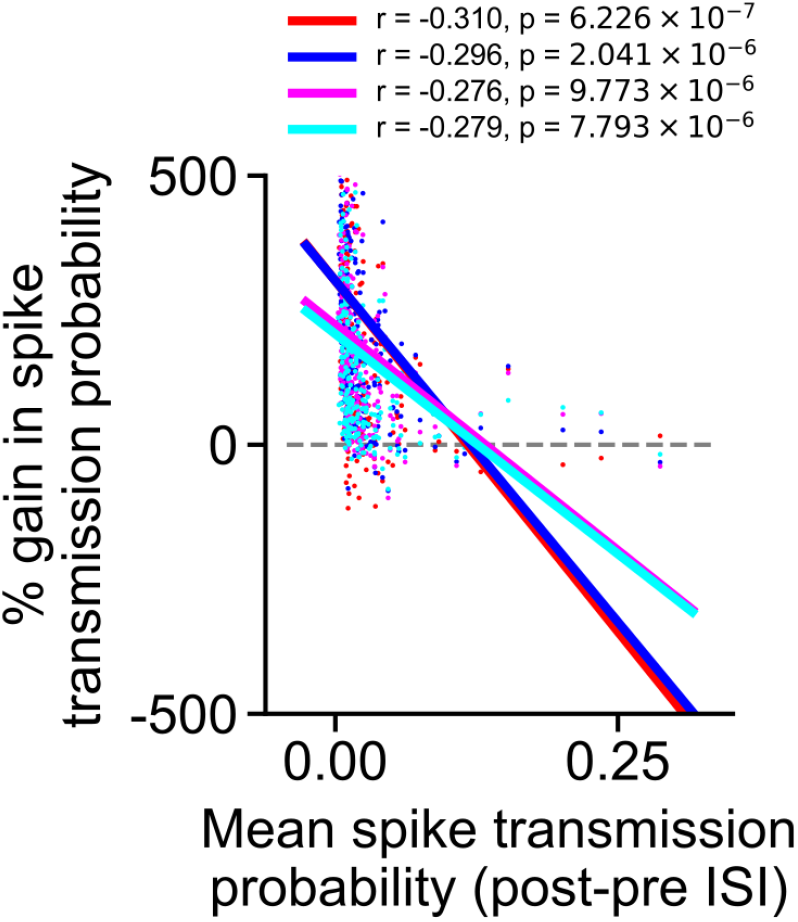
Percentage gain in spike transmission vs. average spike transmission. The percentage gain in spike transmission probability of the post-pre ISIs is negatively correlated to the average post-pre ISI spike transmission probability, with longer ISI ranges having less negative correlation (*r* = -0.310, -0.296, -0.276, -0.279, *p* = 6.23 x 10^-7^, 2.04 x 10^-6^, 9.77 x 10^-6^, 7.79 x 10^-6^ for ISI ranges of 5-25, 25-45, 45-65, and 65-85 ms, *n* = 249 pairs, *n* = 168 RGCs, *n* = 19 experiments, *n* = 18 mice). Red, blue, magenta, and cyan correspond to ISI ranges of 5-25, 25-45, 45-65, and 65-85 ms, respectively.

**Figure S7:**
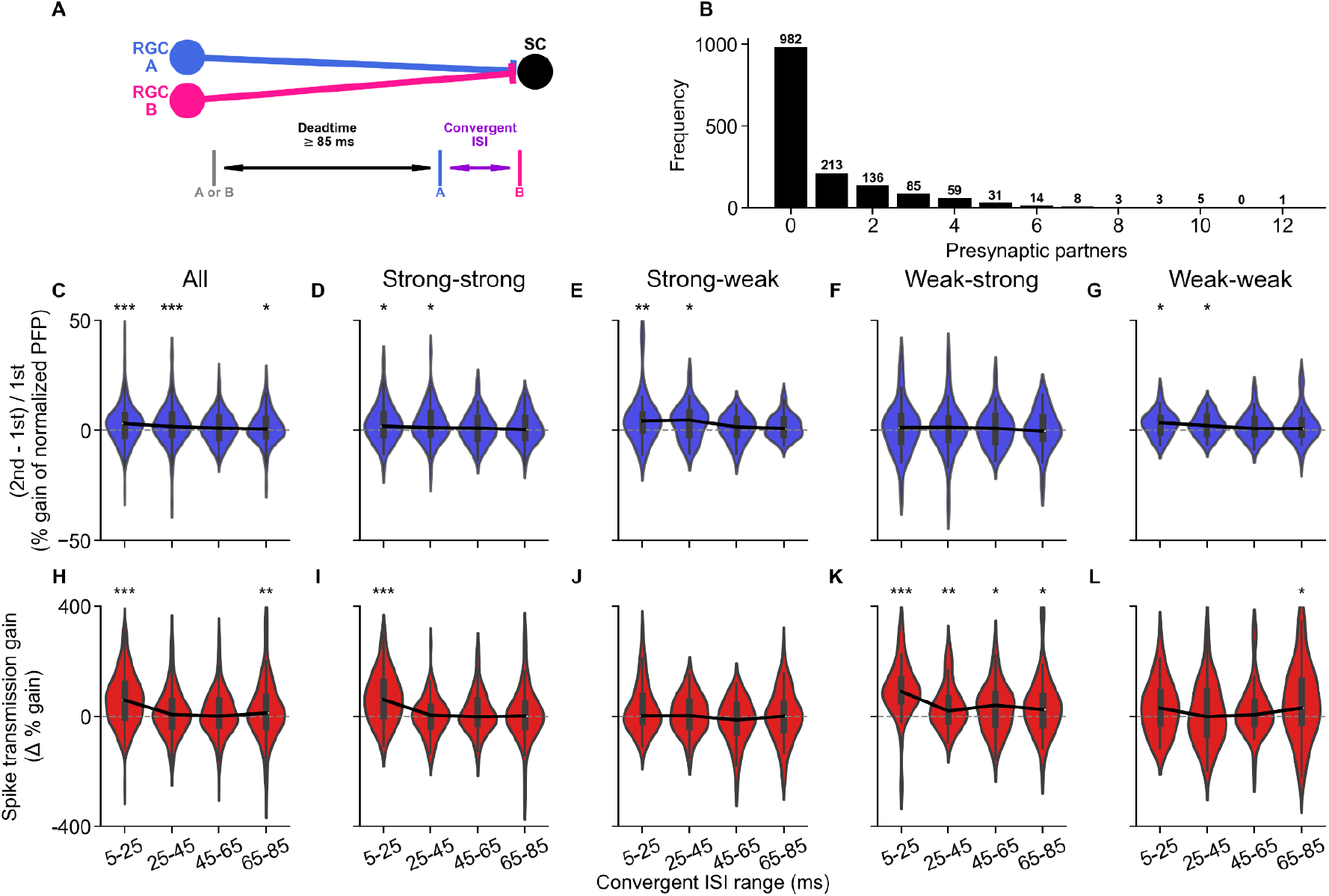
Convergent inputs facilitate the spike transmission probability more than PFP amplitude. (A) Scheme showing the convergent inputs from different presynaptic RGCs to a same postsynaptic SC neuron. The ISI between the paired convergent inputs need to have at least 85 ms dead time in order to be considered in the analyses. (B) The distribution of the number of presynaptic partners for the postsynaptic SC neurons. (C-G) The percentage of the normalized second PFP amplitude over the normalized first PFP amplitude (see Methods). The presynaptic RGC is considered as strong if the average spike transmission to the postsynaptic SC neuron is ≥0.02, otherwise it is considered as weak connection. There is only a very small facilitation for all pairs (C; Q1: -2.52%, -1.96%, -3.52%, -2.63%; median: 3.07%, 1.78%, 1.02%, 0.61%; Q3: 7.1%, 7.35%, 5.75%, 5.3%; *p* = 6.63 x 10^-5^, 1.25 x 10^-4^, 0.075, 0.036 for ISI ranges of 5-25, 25-45, 45-65, and 65-85 ms, two-sided Wilcoxon signed-rank test for first PFP vs. second PFP, *n* = 162 RGC pairs, *n* = 11 experiments, *n* = 11 mice), which is mainly stemming from the weak second spike of strong-weak RGC pairs (E; Q1: 0.03%, -2.04%, -1.83%, -2.04%; median: 4.31%, 4.66%, 1.6%, 0.83%; Q3: 7.49%, 8.3%, 5.31%, 5.01%; *p* = 4.12 x 10^-3^, 0.015, 0.279, 0.08; *n* = 36 RGC pairs). The strong-strong (D; Q1: -2.11%, -1.66%, -3.63%, -3.29%; median: 1.94%, 1.24%, 1%, 0.38%; Q3: 7.39%, 7.97%, 5.62%, 5.59%; *p* = 0.013, 0.011, 0.369, 0.265; *n* = 66 RGC pairs), weak-strong (F; Q1: -4.98%, -4.03%, -5.05%, -3.75%; median: 1.33%, 1.43%, 0.91%, -0.32%; Q3: 6.57%, 5.55%, 6.66%, 5.98%; *p* = 0.698, 0.612, 0.84, 0.826; *n* = 34 RGC pairs), and weak-weak (G; Q1: -0.83%, -1.19%, -1.53%, -1.81%; median: 3.46%, 2.17%, 0.81%, 0.8%; Q3: 5.68%, 6.34%, 5.1%, 4.52%; *p* = 0.018, 0.036, 0.237, 0.173; *n* = 26 RGC pairs) RGC pairs have little to no PFP facilitation. (H-L) The percentage difference in spike transmission probability gain between the first and second spikes from different RGCs (see Methods). There is a facilitation for all pairs (H; Q1: -3.94%, -36.75%, -33.68%, -38.24%; median: 58.79%, 7.11%, 0.53%, 12.28%; Q3: 119.14%, 55.44%, 58.3%, 70.76%; *p* = 3.47 x 10^-17^, 0.072, 0.086, 0.008 for ISI ranges of 5-25, 25-45, 45-65, and 65-85 ms, two-sided Wilcoxon signed-rank test for first spike vs. second spike, *n* = 228 connection pairs, *n* = 162 RGC pairs, *n* = 11 experiments, *n* = 11 mice), which is mainly stemming from the pairs that have a second spike from a strong RGC connection, namely strong-strong (I; Q1: 0.96%, -35.83%, -27.36%, -38.22%; median: 60.56%, 4.13%, -1.73%, 1.16%; Q3: 124.6%, 36.39%, 58.64%, 46.18%; *p* = 1.15 x 10^-8^, 0.991, 0.442, 0.574; *n* = 86 connection pairs) and weak-strong (K; Q1: 55.31%, -16.24%, -30.88%, -31.55%; median: 90.33%, 19.2%, 40.43%, 24.71%; Q3: 134.24%, 66.32%, 82.59%, 73.01%; *p* = 2.44 x 10^-9^, 0.004, 0.02, 0.019; *n* = 58 connection pairs) RGC pairs. There is no obvious facilitation in the strong-weak (J; Q1: -20.85%, -38.32%, -58.03%, -48.92%; median: 2.24%, 2.58%, -13.74%, 0.7%; Q3: 72.73%, 54.75%, 41.3%, 45.16%; *p* = 0.181, 0.429, 0.232, 0.88; *n* = 46 connection pairs) and weak-weak (L; Q1: -26.14%, -65.5%, -27.37%, -24.65%; median: 30.02%, -1.3%, 5.98%, 29.74%; Q3: 89.71%, 94.08%, 52.38%, 128.82%; *p* = 0.099, 0.875, 0.321, 0.027; *n* = 38 connection pairs) RGC pairs.

**Figure S8:**
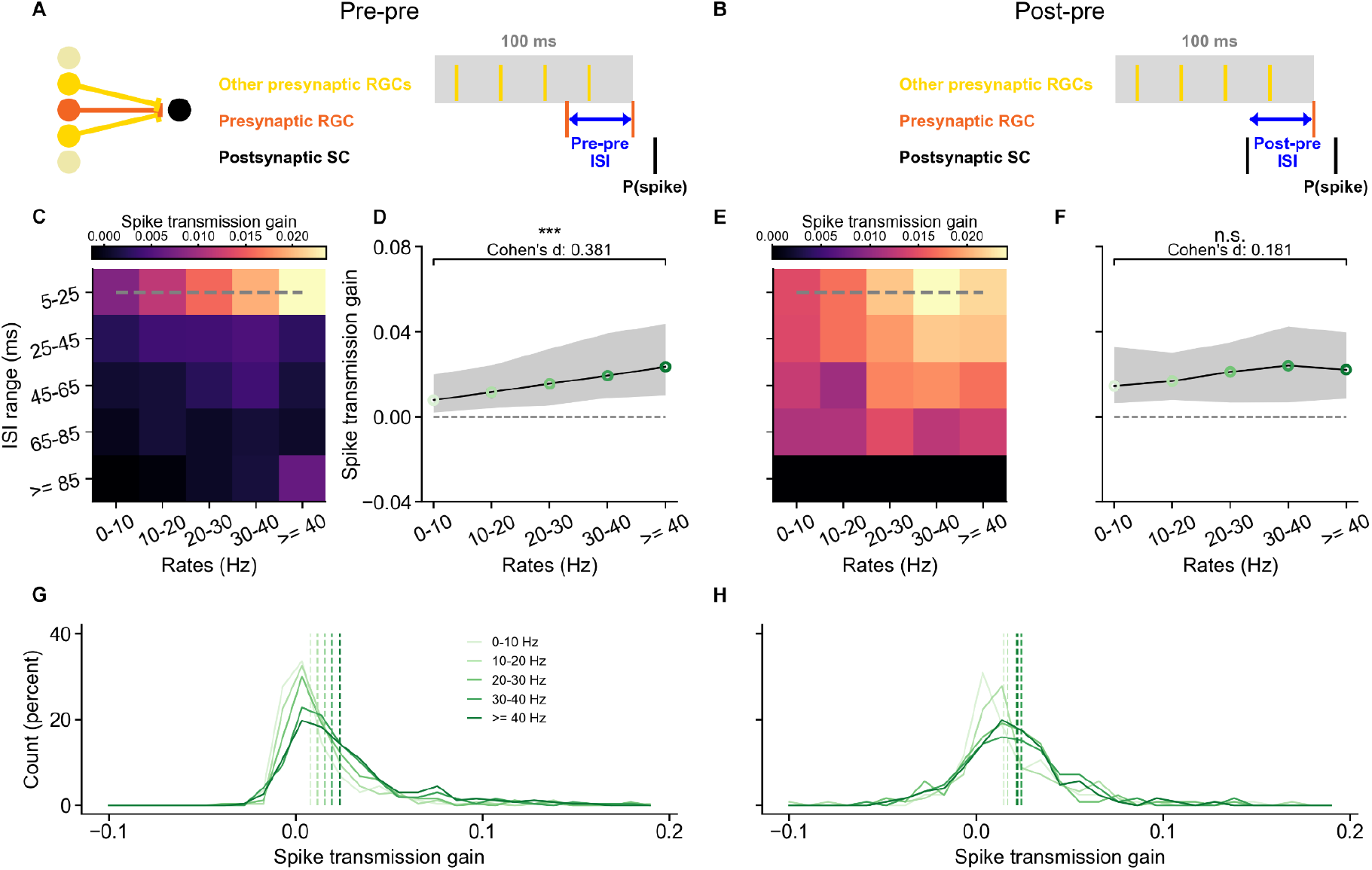
Effects of input rates on short-term facilitation. (A-B) Schematics showing the input-rate window used for the pre-pre (A) and post-pre (B) ISI conditions. (C and E) The median raw spike transmission gain of different pre-pre (C) and post-pre (E) ISIs for different input-rate levels. (D and F) The raw spike transmission gain of 5-25 ms ISI against different input rates for the pre-pre (D; 0-10 Hz vs. ≥40 Hz: *p* = 6.75 x 10^-16^; for input rates of 0-10, 10-20, 20-30, 30-40, and ≥40 Hz, Q1: 0.00214, 0.00407, 0.00553, 0.00889, 0.0102; median: 0.00793, 0.0117, 0.0157, 0.0194, 0.0236; Q3: 0.02, 0.0242, 0.032, 0.0393, 0.0438, *n* = 271 pairs, *n* = 163 RGCs, *n* = 17 experiments, *n* = 16 mice) and post-pre (F; 0-10 Hz vs. ≥40 Hz: *p* = 0.099; for input rates of 0-10, 10-20, 20-30, 30-40, and ≥40 Hz, Q1: 0.00652, 0.00788, 0.00683, 0.00688, 0.00857; median: 0.0146, 0.0169, 0.0212, 0.0241, 0.0222; Q3: 0.0329, 0.0302, 0.035, 0.0426, 0.0397, *n* = 126 pairs, *n* = 97 RGCs, *n* = 15 experiments, *n* = 14 mice) ISI conditions. The black line indicates the median and the shaded region indicates the Q1 and Q3 of the raw spike transmission gain. (G-H) Distributions of raw gain of spike transmission probability for different input rates at 5-25 ms pre-pre (G) and post-pre (H) ISIs. The dashed lines indicate the median of the distributions.

**Figure S9:**
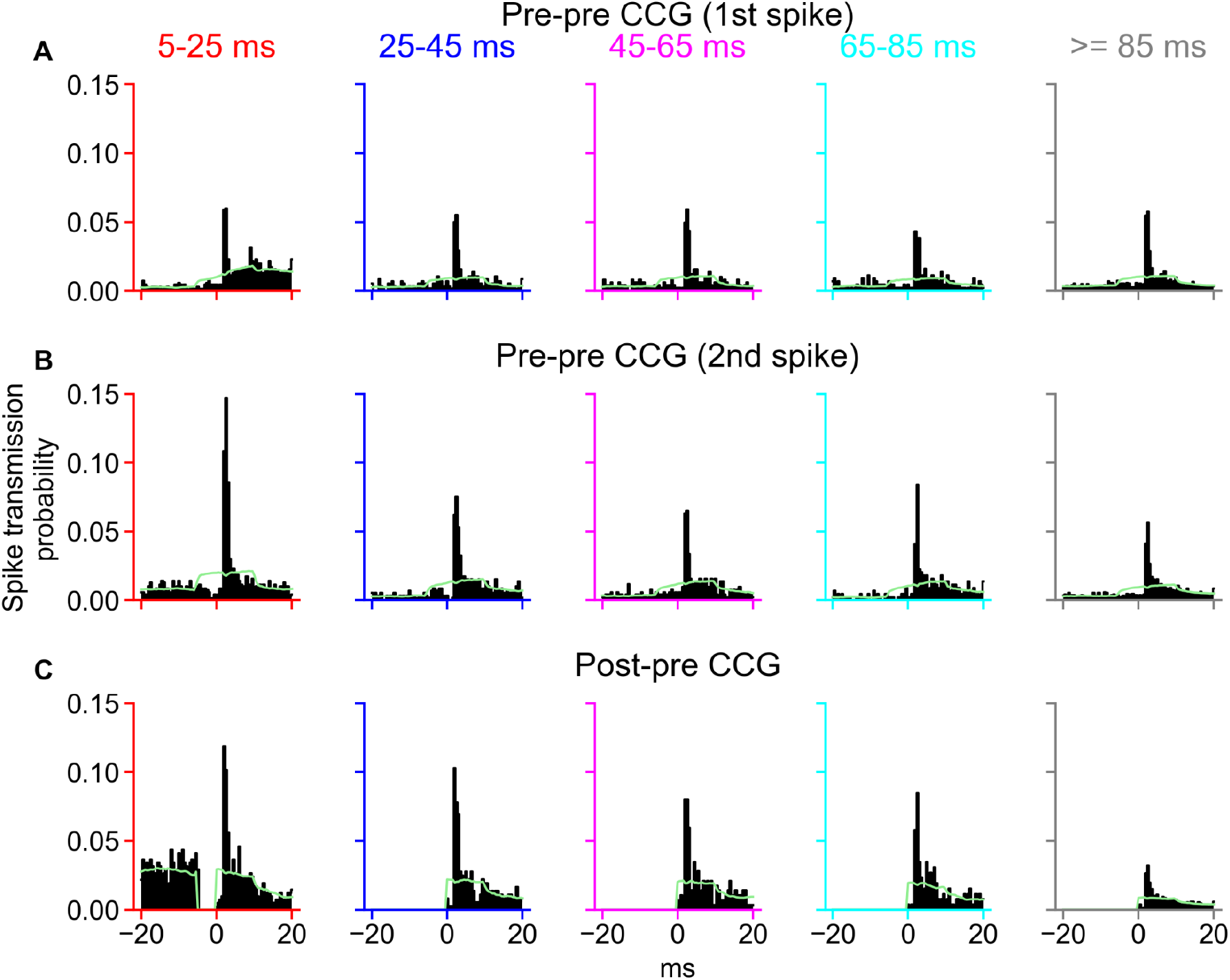
Model CCGs of one presynaptic neuron and the postsynaptic neuron for the pre-pre and post-pre ISI conditions. (A) The CCGs of the first spike for the pre-pre ISI condition. (B) The CCGs of the second spike for the pre-pre ISI condition. (C) The CCGs for the post-pre ISI condition. The green line indicates the baseline of the CCG.

